# AKT-mediated phosphorylation of TSC2 controls stimulus- and tissue-specific mTORC1 signaling and organ growth

**DOI:** 10.1101/2024.09.23.614519

**Authors:** Yann Cormerais, Samuel C. Lapp, Krystle C. Kalafut, Madi Y. Cissé, Jong Shin, Benjamin Stefadu, Jean Personnaz, Sandra Schrotter, Angelica D’Amore, Emma R. Martin, Catherine L. Salussolia, Mustafa Sahin, Suchithra Menon, Vanessa Byles, Brendan D. Manning

**Author notes:** IRSD, Université de Toulouse, INSERM, INRAE, ENVT, Univ Toulouse III - Paul Sabatier (UPS), Toulouse, France. Cell Signaling Technologies, Inc, Beverly, MA, 01915, USA. Novartis Institutes for BioMedical Research, Cambridge, MA, 02139, USA. These authors contributed equally (Samuel C. Lapp, Krystle C. Kalafut, Madi Y. Cissé, Jong Shin).

## Abstract

Mechanistic target of rapamycin (mTOR) complex 1 (mTORC1) integrates diverse intracellular and extracellular growth signals to regulate cell and tissue growth. How the molecular mechanisms regulating mTORC1 signaling established through biochemical and cell biological studies function under physiological states in specific mammalian tissues are unknown. Here, we characterize a genetic mouse model lacking the 5 phosphorylation sites on the tuberous sclerosis complex 2 (TSC2) protein through which the growth factor-stimulated protein kinase AKT can active mTORC1 signaling in cell culture models. These phospho-mutant mice (TSC2-5A) are developmentally normal but exhibit reduced body weight and the weight of specific organs, such as brain and skeletal muscle, associated with cell intrinsic decreases in growth factor-stimulated mTORC1 signaling. The TSC2-5A mouse model demonstrates that TSC2 phosphorylation is a primary mechanism of mTORC1 activation in some, but not all, tissues and provides a genetic tool to facilitate studies on the physiological regulation of mTORC1.

---

Cell intrinsic growth factor-stimulated signaling pathways are believed to coordinate organ development, growth, and function in metazoans. While the molecular mechanisms underlying key signaling pathways have been established through previous biochemical and cell biological studies, how these mechanisms function in physiological settings at the organismal and tissue level are poorly understood. Genetic evidence indicates that the PI3K-mTORC1 pathway is a critical determinant of cell and tissue growth, including in rodents and humans. However, how PI3K signals to mTORC1 to influence the physiological growth of mammalian tissues is unknown.

In addition to being central to the control of insulin-responsive glucose metabolism^1,2^, PI3K-AKT signaling can promote anabolic metabolism and tissue growth, in part, through activation of the mechanistic target of rapamycin (mTOR) complex 1 (mTORC1), which has the capacity to integrate signals from multiple additional pathways that can also reflect local and systemic nutrient and energy status^3^. Through cell culture studies, several molecular mechanisms have been proposed for AKT signaling to activate mTORC1 (**Fig-1A**). The core essential components of mTORC1 are the Ser/Thr kinase mTOR, lethal with SEC13 protein 8 (LST8), and regulatory-associated protein of mTORC1 (RAPTOR). Originally, AKT was found to directly phosphorylate mTOR on S2448^4–6^, a phosphorylation event that correlates with increased mTORC1 signaling. However, a molecular effect of S2448 phosphorylation on mTORC1 signaling has yet to be found, and it can occur on mTOR within both mTORC1 and a second mTOR-containing complex, mTORC2, which functions upstream of AKT^6,7^. Furthermore, mTOR-S2448 is now believed to be primarily phosphorylated by S6K1, an AKT-related protein kinase that is activated downstream of mTORC1^8,9^. AKT also directly phosphorylates a dispensable component of mTORC1 dubbed proline-rich AKT substrate of 40 kD (PRAS40)^10^, which can associate with mTORC1 in an inhibitory manner but is also a direct substrate of mTORC1^11–15^. AKT-mediated phosphorylation of T246 on PRAS40 has been suggested to relieve an inhibitory effect of PRAS40 on mTORC1, thereby contributing to mTORC1 activation downstream of AKT^13,14^.

**Figure 1:**
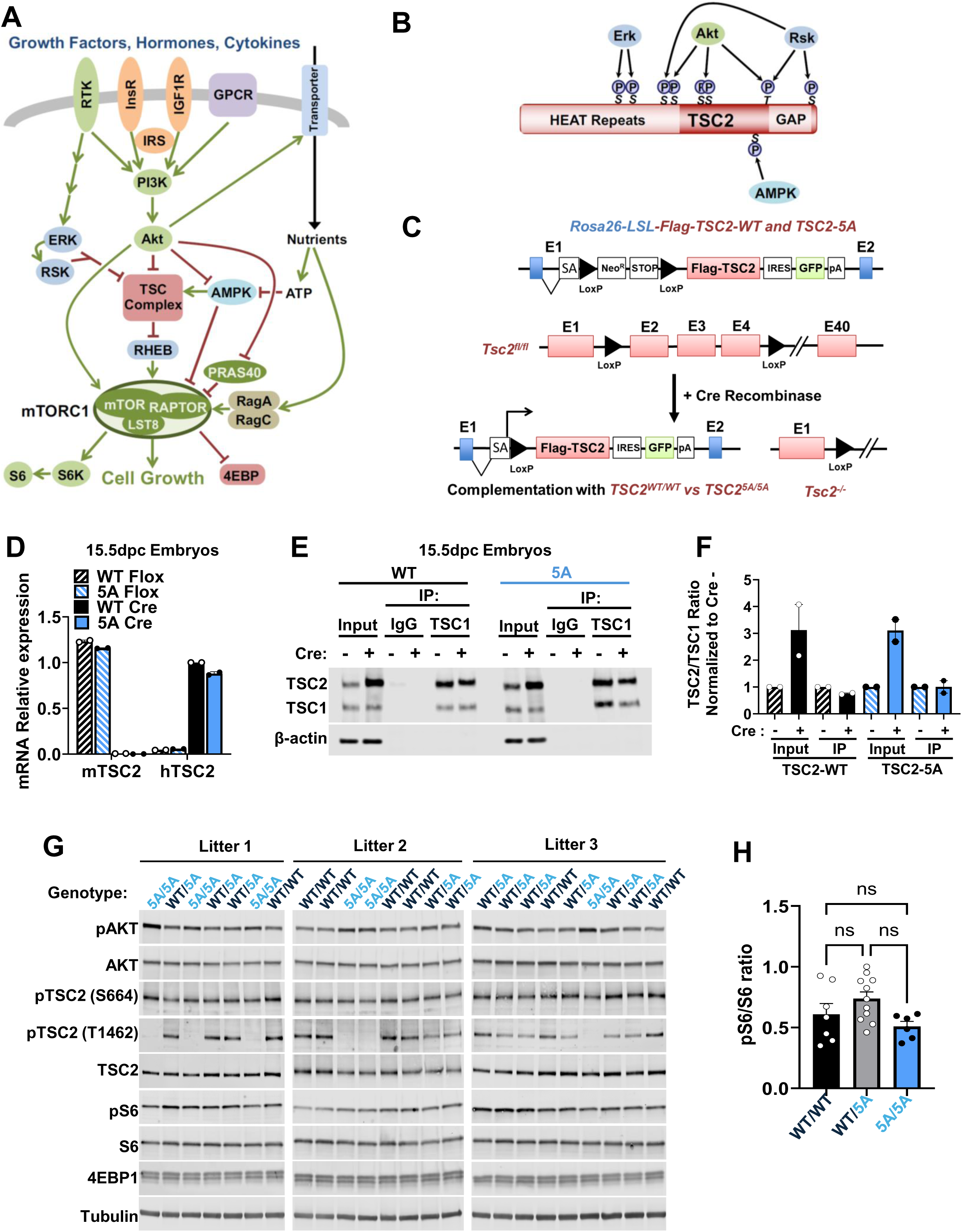
A genetic mouse model to disconnect AKT signaling from TSC2 regulation. (A) Schematic of proposed mechanisms of mTORC1 regulation. (B) Schematic of the domain structure and phosphorylation sites on TSC2. (C) Schematic of the strategy for generating conditional TSC2-WT and −5A mice, involving transgene knock-ins at the *Rosa26* locus and a *Tsc2*-floxed allele. (D) qPCR quantification of mRNA expression of the mouse and human TSC2 transcripts in *Rosa26-LSL-TSC2-WT; Tsc2^fl/fl^* (WT Flox) or *Rosa26-LSL-TSC2-5A; Tsc2^fl/fl^* (5A Flox) and following expression of CMV-Cre, *Rosa26-TSC2-WT; Tsc2^−/−^* (WT Cre) or *Rosa26-TSC2-5A; Tsc2^−/−^* (5A Cre) 15.5 dpc whole embryos. Data are graphed as mean ± SEM; N = 2 biological replicates per genotype. (E) Immunoblots of total protein (Input) and immunoprecipitations (IP) with TSC1 or IgG control antibodies from protein extracts from *Rosa26-LSL-TSC2-WT* or *-5A; Tsc2^fl/fl^* (Cre -) and *Rosa26-TSC2-WT* or *-5A; Tsc2^−/−^* (Cre +) 15.5 dpc whole embryos are shown. (F) Quantification of the TSC2/TSC1 ratio observed in (E), graphed as mean fold change from Cre-control ± SEM; N = 2 biological replicates per genotype. (G) Immunoblots of protein extracts from 15.5 dpc whole embryos obtained from 3 independent litters from heterozygous *Rosa26-TSC2^WT/5A;^ Tsc2^−/−^* crosses are shown. (H) Quantification of pS6/S6 ratios from the data in (G), graphed as mean ± SEM; N = 7 TSC2-WT/WT, 11 TSC2-WT/5A, and 6 TSC2-5A/5A. Statistical analysis in (H) was performed using the Kruskal-Wallis test: ns (not significant), p > 0.5.

mTORC1 is activated through a ubiquitously expressed small GTPase named RAS-homolog enriched in brain (RHEB), which in its GTP-bound state directly engages and activates mTOR within mTORC1^3^. The only known regulator of RHEB is the tuberous sclerosis complex (TSC) protein complex, which is comprised of the TSC1, TSC2, and TBC1D7 proteins and acts as a GTPase-activating protein (GAP) for RHEB through a C-terminal domain on TSC2. The GAP activity of the TSC complex maintains RHEB in its GDP-bound state, thereby serving as an essential negative regulator of mTORC1 signaling. Importantly, AKT can promote the RHEB-mediated activation of mTORC1 by phosphorylating 5 residues on TSC2 (human full-length numbering: S939, S981, S1130, S1132, T1462) within the TSC complex, thereby relieving its inhibition of RHEB (**Fig-1A,B**)^16–18^. Other growth factor signaling pathways, including the ERK-RSK pathway, can also activate mTORC1 via phosphorylation of both distinct and overlapping residues on TSC2^19,20^. Illustrating the importance of the TSC complex in suppressing mTORC1 activation, genetic loss of TSC complex components result in constitutive, growth-factor independent activation of mTORC1 signaling and an increase in cell size^21–24^.

Intracellular energy and nutrients also play an essential role in the regulation of mTORC1. While there are multiple mechanisms through which mTORC1 can sense even subtle decreases in intracellular ATP^25^, the initial mechanisms involve the AMP-activated protein kinase (AMPK). AMPK phosphorylates and activates TSC2 within the TSC complex while also phosphorylating RAPTOR within mTORC1, both events serving to inhibit mTORC1 in response to energy depletion (**Fig-1A,B**)^26–28^. In some settings, AKT can phosphorylate and inhibit AMPK^29^, suggesting yet another possible route from AKT to enhanced mTORC1 signaling. mTORC1 activation is also exquisitely sensitive to the depletion of specific nutrients, including amino acids and glucose. A major mechanism by which mTORC1 senses these nutrients is through the RAG family of GTPases, which under nutrient-replete conditions are in the proper GTP/GDP-bound state to directly bind mTORC1 and dock the complex to the surface of the lysosome^30–35^. Unlike RHEB, the RAG GTPases do not activate mTORC1 but bring it to the lysosome where it can be activated by a subpopulation of RHEB that is present there. As AKT signaling can promote glucose and amino acid uptake into cells^36^, it could, theoretically, also influence the RAG pathway to indirectly promote mTORC1 activation (**Fig-1A**). Thus, AKT signaling appears to influence the activation state of mTORC1 in cells through multiple mechanisms, the importance of which have yet to be established *in vivo*.

While the physiological functions of the specific signaling mechanisms described above are unknown, several mouse models have been generated to help define the general role of the core mTORC1 signaling components. Germline knockout of *Tsc1*, *Tsc2*, or *Rheb*, encoding the key mTORC1-proximal components involved in growth factor signaling to mTORC1, or genes encoding components of mTORC1 itself, such as Mtor, Mlst8, or Rptor, result in embryonic lethality at various early stages of development, thus preventing a broader ability to study the role of these proteins in organismal growth and physiology^22,37–41^. Homozygous knockout of the third core component of the TSC complex, TBC1D7, does not influence viability but results in increases in brain and muscle fiber size, illustrating the sensitivity of these tissues to even modest changes in mTORC1 signaling^42^. Several conditional deletion models have been used to study the tissue-specific consequences of either complete constitutive activation of mTORC1 (*Tsc1* or *Tsc2* knockouts)^43–48^ or complete loss of mTORC1 activity (*Rheb* or *Rptor* knockouts)^49–53^.

While these models are helpful to understand the consequences of extreme, all or nothing, effects on mTORC1 signaling, the highly integrative nature of mTORC1 regulation is lost and the phenotypes arising are often secondary to ablation of these essential signaling components. For instance, chronically elevated mTORC1 signaling from loss of the TSC complex in specific tissues can yield an initial increase in tissue size but ultimately results in proteotoxic stress and tissue damage with age and is accompanied by a strong feedback inhibition of PI3K-AKT signaling^44,45,48,54,55^, effects that do not necessarily reflect physiological consequences of regulated mTORC1 signaling.

Here, we describe the generation and initial characterization of a novel genetic mouse model that conditionally replaces endogenous TSC2 with a phospho-mutant of TSC2 that cannot be phosphorylated and regulated by AKT (TSC2-5A), thus eliminating a major mode of AKT signaling to mTORC1. We find that TSC2-5A embryos have normal mTORC1 signaling and are born at mendelian ratios, indicating that this regulation is dispensable for mouse development. However, primary cells derived from these embryos and mice display defects in both basal and growth factor-stimulated mTORC1 signaling, and TSC2-5A mice exhibit a decrease in lean body mass and the weight of specific organs relative to their TSC2-WT counterparts. We characterize the brain and skeletal muscle phenotypes and find cell intrinsic changes in mTORC1 signaling and function accompanying these differences in organ size. Thus, this study demonstrates the physiological importance of AKT-mediated phosphorylation of TSC2 for the tissue-specific control of mTORC1 signaling and organ growth.

## Results

### A genetic mouse model to disconnect AKT signaling from TSC2 regulation

Support for the several previously proposed mechanisms of AKT-mediated activation of mTORC1, discussed above, stems largely from studies performed in immortalized proliferating cell lines. To begin to understand the physiological relevance of AKT-dependent regulation of mTORC1 by growth factor signaling, we examined this regulation in non-proliferative, terminally differentiated cells. Primary hepatocytes, myotubes and neurons were isolated from C57BL/6J mice and stimulated with insulin or insulin-like growth factor 1 (IGF1) in the presence or absence of the AKT inhibitor MK2206, the MEK inhibitor trametinib, or the mTORC1 inhibitor rapamycin. While the level of basal mTORC1 signaling varied between these primary cells, insulin/IGF1 acutely stimulated mTORC1 in all cell types, as scored by phosphorylation of the direct mTORC1 substrates S6K1 and 4EBP1 (indicated by immunoblot mobility shift) (**Fig-S1A-C**). In all cell types, this stimulated activity was entirely dependent on AKT, but not MEK, and correlated with the phosphorylation of TSC2 and PRAS40, but not mTOR, on established AKT sites. As expected, rapamycin inhibited mTORC1 signaling to levels below the unstimulated state. These results confirm that AKT is the critical growth factor-stimulated kinase regulating mTORC1 in these primary cells in response to insulin/IGF1 and that multiple mechanisms previously described for the AKT-mediated activation of mTORC1 can be simultaneously activated in such cells.

To critically define the role of the TSC complex as a downstream target of AKT in the control of tissue and organismal mTORC1 signaling, we generated two novel knock-in mouse models conditionally expressing either wild-type TSC2 (TSC2-WT) or phospho-mutant TSC2 containing alanine mutations at the five residues directly phosphorylated by AKT (TSC2-5A). Flag-tagged human TSC2 cDNA was inserted into the Rosa26 locus using the previously described targeting vector Stop-EGFP-Rosa TV^56^, which encodes transcriptional and translational stop sequences flanked by loxP sites (Lox-Stop-Lox or LSL cassette), allowing for temporal and spatial control of TSC2 expression from the ubiquitous Rosa26 promoter upon Cre recombinase expression (**Fig-1C**). Proper 5’ and 3’ insertions of the transgene into the Rosa26 locus were confirmed using PCR (**Fig-S1D,E**). Mice containing the Rosa26-LSL-TSC2-WT or - 5A insertion were crossed into a *Tsc2^fl/fl^*line, where exons 2 through 4 of the endogenous mouse *Tsc2* locus are flanked by loxP sites^42^, allowing coordinated deletion of the endogenous gene and expression of the TSC2 transgenes to entirely replace endogenous TSC2 in a given tissue upon expression of CRE recombinase (**Fig-1C**). In this study, we describe the effects of whole-body allele replacement by crossing the mice into a strain expressing Cre from the ubiquitous cytomegalovirus (CMV) promoter^57^.

We confirmed that the CMV-Cre led to simultaneous whole-body deletion of the endogenous mouse *Tsc2* gene and complementation with either the human TSC2-WT or −5A transgene using qPCR (**Fig-1D**) and PCR (**Fig-S1F,G**) analyses of whole embryos. Importantly, while higher expression of total TSC2 protein was observed in the transgene-activated embryos (**Fig-1E,F and Fig-S1H**), there were no changes in TSC1 abundance and very similar levels of assembled TSC complexes between transgene-expressing (Cre^+^) and non-expressing (Cre^-^) mice and no difference between TSC2-WT and TSC2-5A mice (**Fig-1E,F**). This finding is consistent with the fact that free pools of TSC2 exist in cells and that TSC1 sets the overall level of TSC complex, which is the functional Rheb-GAP^18^. Somewhat surprisingly, we found that ubiquitous expression of either the TSC2-WT or TSC2-5A transgene fully complements the early embryonic lethality of *Tsc2^−/−^* mice^41^, with no overt developmental phenotype. Indeed, heterozygous crosses between *Rosa26-TSC2^WT/5A^;Tsc2^−/−^*mice yielded offspring genotypes in the expected F1 mendelian ratios (**Table 1**). Consistent with this rescue of lethality, normal whole embryo AKT and mTORC1 signaling, as revealed by AKT and S6 phosphorylation, respectively, was detected across three litters from these crosses, with genotyping done in parallel (**Fig-1G,H**). As expected, phosphorylation of TSC2 on the AKT site T1462 was absent specifically in TSC2-5A embryos, but there was no difference in TSC2 phosphorylation at the ERK site S664^19^ between TSC2-WT and −5A embryos (**Fig-1G**). These results demonstrate that the five AKT-mediated phosphorylation sites on TSC2 are dispensable for normal mammalian development, similar to that described in *Drosophila*^58,59^.

**Table1:**
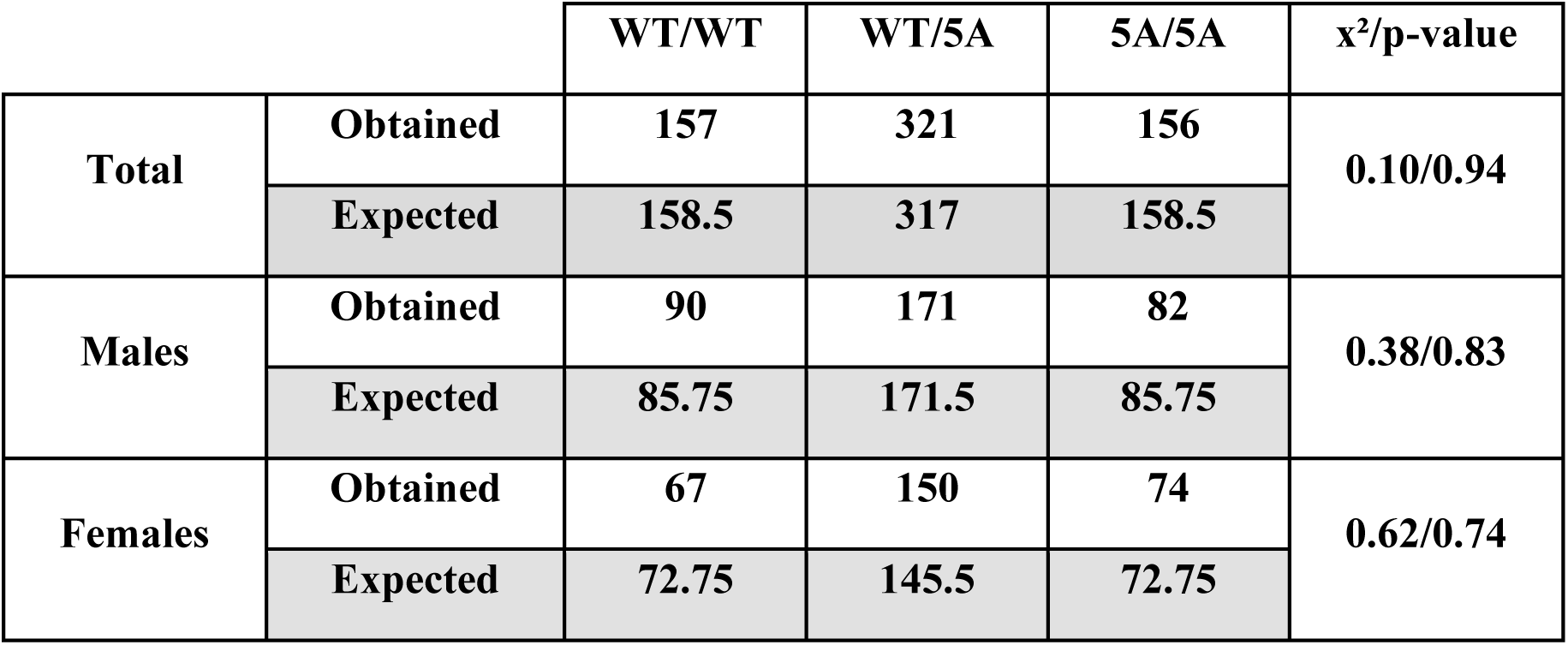
TSC2-5A mice are born at mendelian ratios. Genotypes of 634 offspring of crosses between *Rosa26-TSC2^WT/5A^;Tsc2^−/−^* mice Statistical analysis Chi squared (x²) test.

| | | WT/WT | WT/5A | 5A/5A | $\chi^2$/p-value |
| --- | --- | --- | --- | --- | --- |
| Total | Obtained | 157 | 321 | 156 | 0.10/0.94 |
|  | Expected | 158.5 | 317 | 158.5 |  |
| Males | Obtained | 90 | 171 | 82 | 0.38/0.83 |
|  | Expected | 85.75 | 171.5 | 85.75 |  |
| Females | Obtained | 67 | 150 | 74 | 0.62/0.74 |
|  | Expected | 72.75 | 145.5 | 72.75 |  |

### TSC2-5A cells have decreased AKT-stimulated mTORC1 signaling

To further investigate the role of AKT-mediated phosphorylation of TSC2 in the dynamic regulation of mTORC1 signaling, we isolated three primary mouse embryonic fibroblast (MEFs) lines for each genotype from littermates derived from heterozygous crosses between *Rosa26-TSC2^WT/5A^;Tsc2^−/−^* mice, with all quantifications representing the average of the three biologically independent MEFs lines. TSC2-WT and −5A cells displayed indistinguishable S6K1 phosphorylation (**Fig-2A,B**), cell size (**Fig-2C,D**), and rate of proliferation (**Fig-2E**) under standard nutrient-replete, serum-fed cell culture conditions. However, acute signaling studies revealed a significant general decrease in mTORC1 activation by exogenous growth factors in the TSC2-5A cells. Despite similar AKT activation, TSC2-5A MEFs exhibit a significant decrease in S6K1 phosphorylation compared to TSC2-WT cells in response to acute stimulation with serum, epidermal growth factor (EGF), and insulin (**Fig 2F-H**). This defect in mTORC1 activation was independent of the stimulus concentration used, as the decrease in the TSC2-5A cells was retained even when using a supraphysiological level of insulin (1 μM) (**Fig-2I**). Interestingly, TSC2-5A cells exhibit a 40% reduction in S6K1 phosphorylation compared to TSC2-WT cells even at baseline, following serum starvation (**Fig 2F-I**). We hypothesized that this difference is due to residual AKT signaling to mTORC1 following growth factor withdrawal and that this signal involves AKT-mediated phosphorylation of TSC2. Indeed, TSC2-WT MEFs displayed basal TSC2 phosphorylation at two AKT sites (S939 and T1462) under serum starvation conditions, which was eliminated with the AKT inhibitor MK2206 but not the MEK inhibitor trametinib, the latter of which did decrease the basal phosphorylation of ERK and the ERK site on TSC2 (S664) (**Fig-S2A**). Importantly, the AKT inhibitor, but not the MEK inhibitor, decreased the basal phosphorylation of S6K1, a decrease that was significant in the TSC2-WT cells but not the TSC2-5A cells (**Fig-S2A,B**), indicating that the basal activation of AKT and its phosphorylation of TSC2 is partially responsible for the basal mTORC1 signaling observed in these primary MEFs.

**Figure 2:**
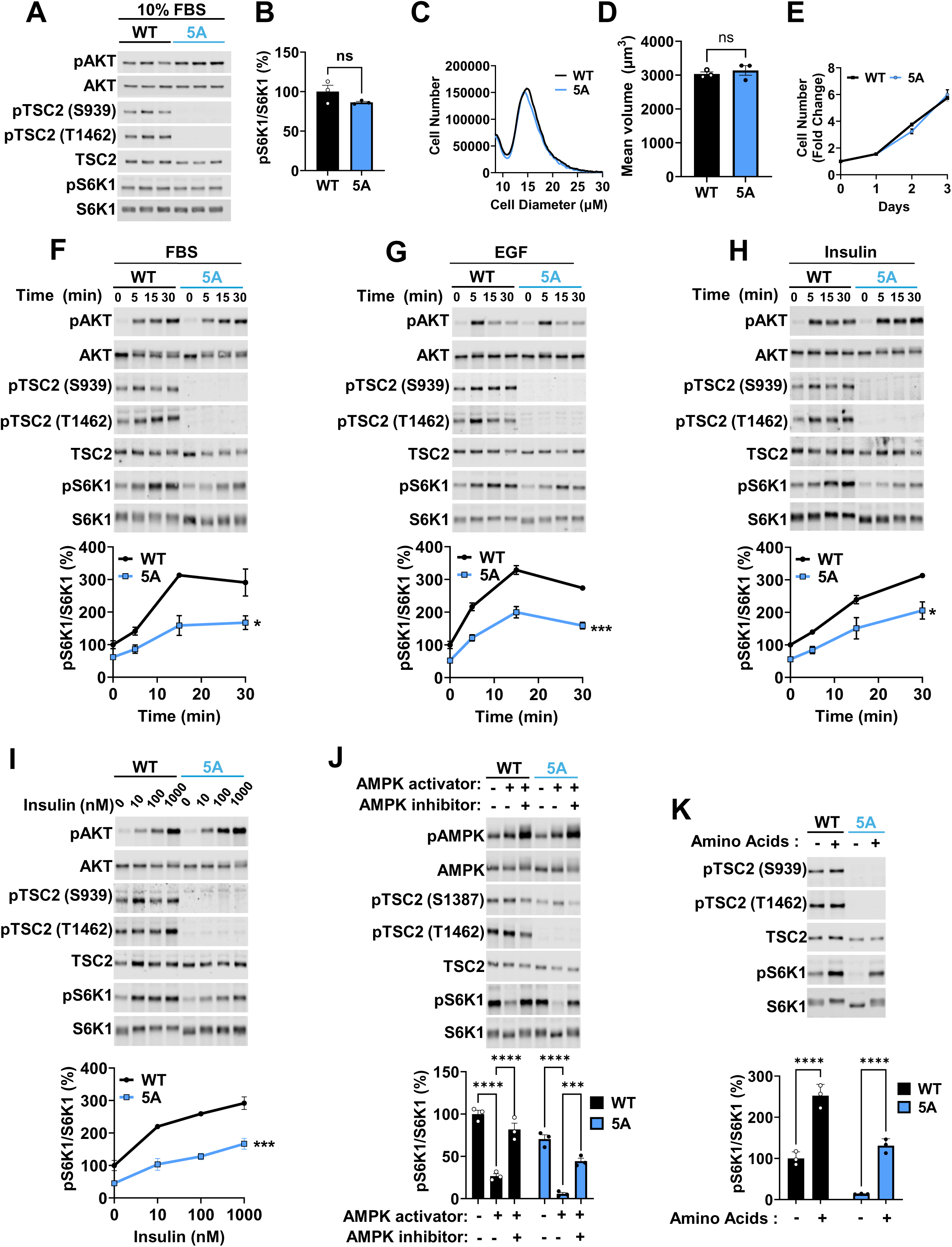
TSC2-5A cells have decreased growth factor-stimulated mTORC1 signaling. (A) Immunoblots of lysates from primary TSC2-WT and TSC2-5A MEFs cultured for 16 h in full serum media (10% FBS). N = 3 biological replicates per genotype. (B) Quantification of pS6K1/S6K1 ratio from (A) expressed as percentage of TSC2-WT MEFs and graphed as mean ± SEM; N = 3 biological replicates per genotype. (C) Representative cell size (diameter) distribution between TSC2-WT and TSC2-5A MEFs cultured as in (A). (D) Mean volume of TSC2-WT and TSC2-5A MEFs cultured as in (A). Data are graphed as mean ± SEM; N = 3 biological replicates per genotype (E) Proliferation curve of the TSC2-WT and TSC2-5A MEFs cultured in full serum media for 3 days. Data are expressed as fold change in cell number relative to day 0 and graphed as mean ± SEM; N = 3 biological replicates per genotype. (F-H) Representative immunoblots and pS6K1/S6K1 ratio quantifications (below) of lysates from primary TSC2-WT and TSC2-5A MEFs serum-starved and then stimulated with time courses of FBS (10%; F), EGF (100 ng/ml; G), or insulin (100 nM; H). Graphed as mean percentage of serum-starved TSC2-WT MEFs ± SEM; N = 3 biological replicates per genotype. (I) Representative immunoblots and pS6K1/S6K1 ratio quantifications (below) of lysates from primary TSC2-WT and TSC2-5A MEFs serum-starved and then stimulated with the indicated concentrations of insulin. Graphed as mean percentage of serum-starved TSC2-WT MEFs ± SEM; N = 3 biological replicates per genotype. (J) Representative immunoblots and pS6K1/S6K1 ratio quantifications (below) of lysates from primary TSC2-WT and TSC2-5A MEFs treated for 1 h with vehicle (-) or an AMPK activator (compound 991, 100 μM) with or without 30 min pretreatment with an AMPK inhibitor (BAY-3827, 1 μM). Graphed as mean percentage of vehicle-treated TSC2-WT MEFs ± SEM; N = 3 biological replicates per genotype. (K) Representative immunoblots and pS6K1/S6K1 ratio quantifications (below) of lysates from primary TSC2-WT and TSC2-5A MEFs deprived of amino acids for 1 h and then, where indicated (+), refed amino acids for 15 min. Graphed as mean percentage of amino acid deprived TSC2-WT MEFs ± SEM; N = 3 biological replicates per genotype. Statistical analysis: (B, D) by Mann-Whitney test; (E-I) repeated measurement two-way ANOVA with Greenhouse–Geisser correction; and (J, K) ordinary two-way ANOVA with Tukey’s multiple comparisons test. *p < 0.05, **p < 0.01, ***p < 0.001, ****p < 0.0001.

To determine the specificity of the model for mTORC1 regulation by growth factors, rather than intracellular energy and nutrients, we compared the effects of AMPK activity and amino acids on mTORC1 signaling between TSC2-WT and −5A cells. When treated with a small molecule allosteric AMPK activator (compound 991^60^), both TSC2-WT and 5A MEFs displayed activating AMPK phosphorylation, TSC2 phosphorylation at the AMPK-regulated site S1387, and decreased S6K1 phosphorylation, with these reciprocal effects on TSC2 and S6K1 phosphorylation blocked by pre-treating cells with an AMPK inhibitor (BAY-3827^61^) (**Fig-2J**). In addition, amino acid starvation and acute refeeding led to a robust activation of mTORC1 in cells of both genotypes (**Fig-2K**). While the fold-induction by amino acids was greater in the TSC2-5A cells, the maximal stimulation was partially reduced relative to TSC2-WT cells. Interestingly, as with growth factor deprivation, the TSC2-5A cells demonstrated greater mTORC1 inhibition in response to both AMPK activation and amino acid starvation (**Fig-2J,K**). Altogether, these results suggest that while dispensable for steady-state activation of mTORC1, the 5 AKT phosphorylation sites on TSC2 are important for the maintenance of basal mTORC1 activity during growth factor or nutrient withdrawal and for acute growth factor-stimulated mTORC1 activation in primary MEFs.

### TSC2-5A mice display reduced body weight and weight of specific organs

While exhibiting no developmental abnormalities, the TSC2-5A male mice weigh significantly less than their TSC2-WT littermates by 8 weeks of age (**Fig-3A**). A similar body weight difference is apparent in female mice albeit after 6 months of age (**Fig-3B**). However, the decreased body weight observed in TSC2-5A mice was not associated with reduced overall body size, as measured by body length (**Fig-3C,D**), but rather was due to a decrease in the weight of several internal organs (**Fig-3E, S3A-C**). Organ weight differences between TSC2-WT and −5A mice were maintained even when normalized to total body weight. This phenotype is temporal in its presentation, with the brain, heart, spleen, and testes being the first organs to show significant reductions in weight in the TSC2-5A male mice at 9 weeks of age (**Fig-S3A**). By 16 weeks, the skeletal muscle, pancreas, and kidneys of TSC2-5A mice also weigh significantly less than those of TSC2-WT mice (**Fig-3E, S3B**). Paradoxically, both the inguinal (iWAT) and epididymal (eWAT) white adipose tissue of TSC2-5A mice weighed more than their WT counterparts, while liver and lung weights were unaffected (**Fig-3E**). In females, the brain and kidneys were the only organs that weighed less in TSC2-5A compared to WT mice at 9 weeks of age (**Fig-S3C**). Together, these results demonstrate that phosphorylation of TSC2 on the 5 AKT-regulated sites promotes postnatal growth in a sex- and organ-specific manner.

**Figure 3:**
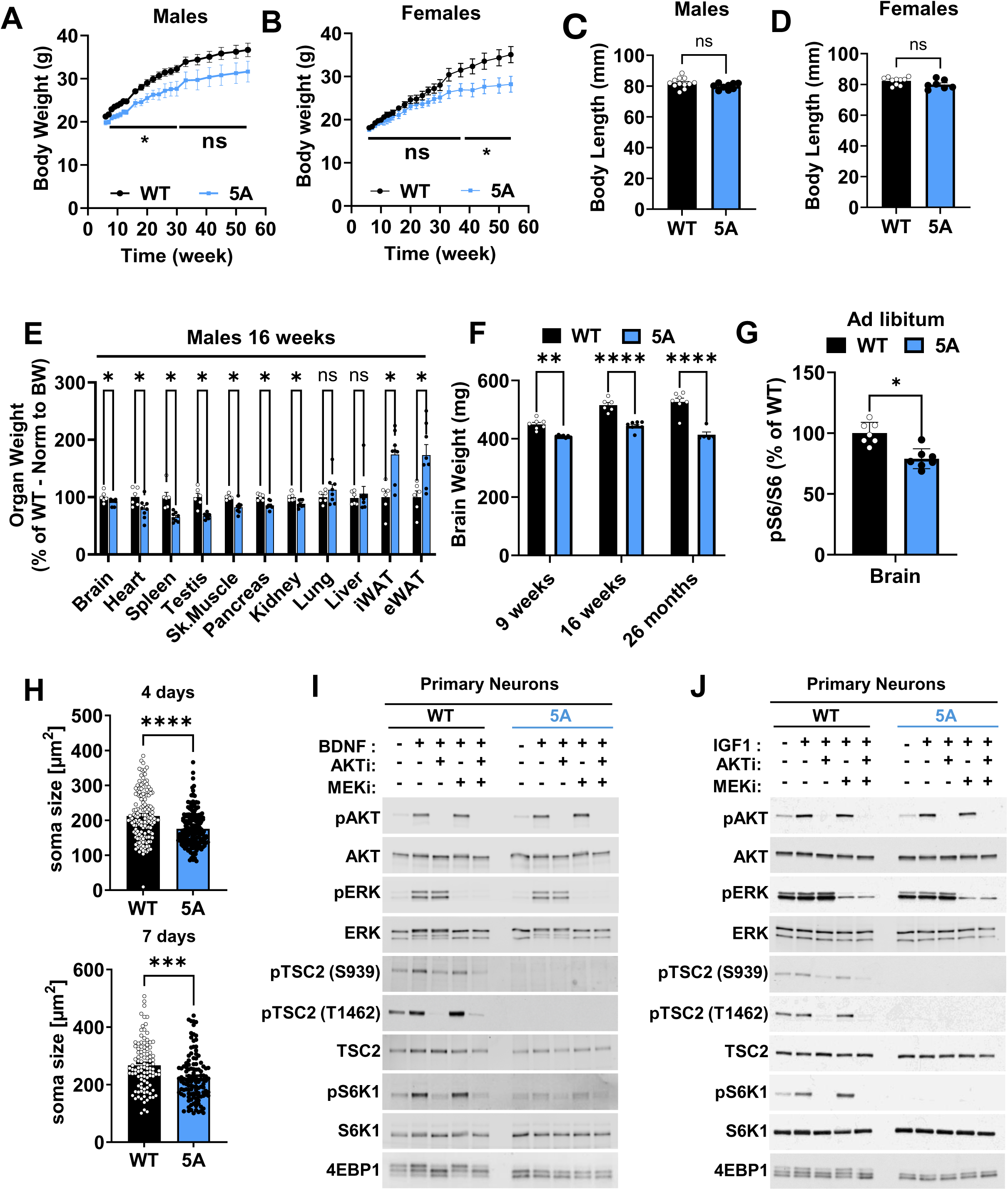
TSC2-5A mice display decreased body and organ-specific weight, including the brain with associated differences in mTORC1 signaling and neuron size. (A) Body weight measured over 12 months in a cohort of TSC2-WT and TSC2-5A littermate males, graphed as mean ± SEM; N = 12 TSC2-WT and 10 TSC2-5A. (B) Body weight measured over 12 months in a cohort of TSC2-WT and TSC2-5A littermate females, graphed as mean ± SEM; N = 10 TSC2-WT and 7 TSC2-5A. (C) Body length of male littermates measured at 11 weeks of age and graphed as mean ± SEM; N = 12 TSC2-WT and 10 TSC2-5A. (D) Body length of female littermates measured at 11 weeks of age and graphed as mean ± SEM; N = 10 TSC2-WT and 7 TSC2-5A. (E) Male organ weights normalized to body weight (BW) measured in 16-week-old TSC2-5A mice, graphed as mean percentage of TSC2-WT littermates ± SEM; N = 6 TSC2-WT and 8 TSC2-5A. (F) Brain weight measurement of male littermates at 9 weeks (N = 7 per genotype), 16 weeks (N = 6 TSC2-WT and 8 TSC2-5A), and 26 months (N = 7 TSC2-WT and 5 TSC2-5A), graphed as mean ± SEM. (G) Ad libitum pS6/S6 ratio measured in brain lysates of 9-week-old TSC2-5A mice (N = 7), graphed as mean percentage of TSC2-WT littermates (N = 7) ± SEM. Quantification of immunoblots in Figure S3D. (H) Soma size (area) of primary hippocampal neurons after 4 and 7 days *in vitro*, graphed as mean ± SEM for 2 independent experiments each. 4 days: N = 149 TSC2-WT and 191 TSC2-5A; 7 days: N = 103 TSC2-WT and 117 TSC2-5A. (I, J) Representative immunoblots of lysates from primary TSC2-WT and TSC2-5A cortical neurons following 4 h of growth factor and amino acid withdrawal, followed by a 15-min stimulation with BDNF (100 ng/mL; I) or IGF1 (100 ng/mL; J) after a 30-min pretreatment with vehicle (-), MK2206 (2 μM, AKTi), trametinib (5 μM, MEKi), or both, as indicated. Statistical analysis: (A, B) by mixed model effect with Šídák’s multiple comparisons; (C-E, G, H) by Mann-Whitney test; and (F) by ordinary two-way ANOVA with Tukey’s multiple comparisons test. *p < 0.05, **p < 0.01, ***p < 0.001, ****p < 0.0001.

### TSC2-5A mice display decreased brain weight, with neuron-intrinsic differences in cell size and mTORC1 signaling

Previous studies using genetic mouse models with increased or decreased mTORC1 signaling in the brain demonstrate that mTORC1 activity promotes brain growth^42,62,63^. Given that the brain was the first organ to exhibit a weight decrease in both male and female TSC2-5A mice relative to TSC2-WT mice (**Fig-S3A,C**) and that this growth defect was sustained out to 26 months of age (**Fig-3F**), we investigated mTORC1 signaling in TSC2-WT and −5A brain tissue and cultured primary neurons. A modest, but significant, 20% decrease in mTORC1 signaling was detected in whole-brain extracts from *ad libitum* fed TSC2-5A mice at 9 weeks of age, as revealed by S6 phosphorylation (**Fig-3G, S3D**). To determine whether these size and signaling changes extend to individual neurons, primary hippocampal and cortical neurons were cultured from litters of TSC2-WT and TSC2-5A embryos. Cultured primary TSC2-5A hippocampal neurons were significantly smaller than their TSC2-WT counterparts after both 4 and 7 days *in vitro* (**Fig-3H**). While both brain-derived neurotrophic factor (BDNF) and IGF1 acutely activated AKT signaling in these primary cortical neurons, BDNF also stimulated ERK signaling, with similar upstream stimulation observed between TSC2-WT and −5A neurons (**Fig-3-I,J**). Despite BDNF stimulating both AKT and ERK, the stimulation of mTORC1 signaling (S6K1 phosphorylation and 4EBP1 mobility shift) by either BDNF or IGF1 was completely blocked with an AKT inhibitor, but not a MEK inhibitor, in TSC2-WT neurons. Consistent with the AKT-dependence of these neuronal growth signals, TSC2-5A neurons exhibited strongly attenuated mTORC1 activation in response to both BDNF and IGF1(**Fig-3I,J**). It is worth noting that we detected a small induction of S6K1 phosphorylation in response to BDNF in the TSC2-5A neurons that remained sensitive to AKT inhibition, suggesting a minor role for parallel AKT-dependent mechanisms in neuronal mTORC1 activation (**Fig-3I**), as detected in primary MEFs. Altogether, these results demonstrate that AKT-mediated phosphorylation of TSC2 is the dominant signal regulating neuronal growth factor-mediated mTORC1 activation and growth and that the AKT-TSC2-mTORC1 signaling circuit regulates brain and neuron size.

### Tissue-specific effects on feeding-induced mTORC1 signaling and organ size in the TSC2-5A mice

AKT and mTORC1 are well established to be activated by insulin signaling, but in our comparison of organ weights between TSC2-WT and TSC2-5A mice two major insulin-responsive tissues had distinct phenotypes, with the skeletal muscle weighing less and the liver exhibiting no change in weight (**Fig-3E**). In contrast to the brain, mTORC1 signaling in skeletal muscle and liver tissue from TSC2-WT and −5A mice was not significantly different under ad libitum fed conditions (**Fig-4A,S4A**). It is worth noting that these tissues exhibit much more variability in mTORC1 signaling under such conditions than does the brain. However, mTORC1 signaling is known to be highly sensitive to fasting and feeding in these tissues, and we found that acute mTORC1 activation by feeding was significantly reduced in skeletal muscle (quadriceps and gastrocnemius) from TSC2-5A mice, despite robust stimulation of AKT activation (**Fig-4B,C**). In contrast, while fasting mTORC1 signaling was significantly lower in liver tissue from TSC2-5A compared to TSC2-WT mice, feeding-induced mTORC1 activation was normal (**Fig-4D**). These results demonstrate that stimulated phosphorylation of TSC2 is required for feeding-induced mTORC1 activation in muscle, but dispensable in the liver, highlighting key differences in mTORC1 regulation in these metabolic tissues.

**Figure 4:**
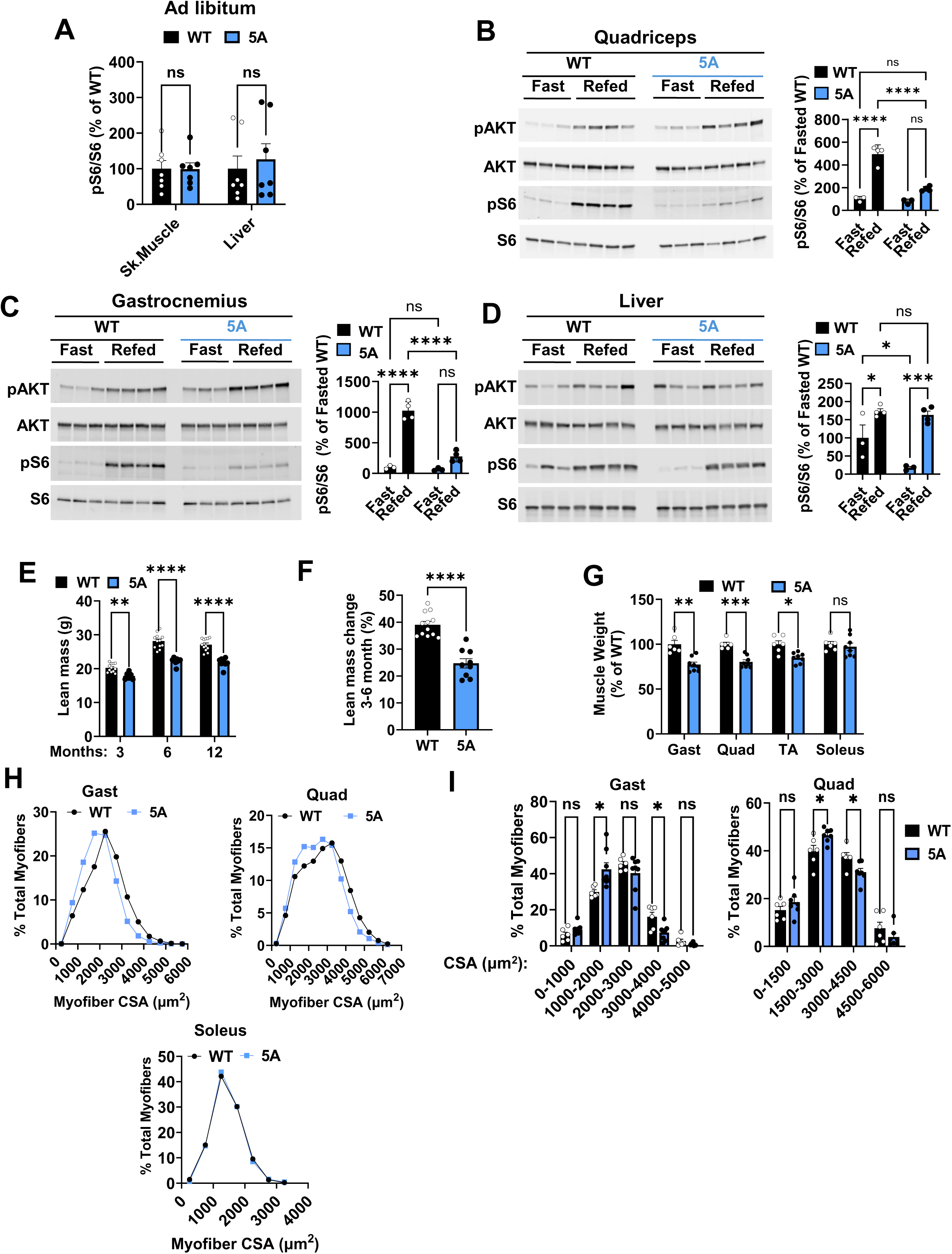
Tissue-specific effects on feeding-induced mTORC1 signaling and organ size in skeletal muscle from TSC2-5A mice. (A) Quantification of male skeletal muscle (gastrocnemius) and hepatic ad libitum pS6/S6 ratios in lysates from 9-week-old TSC2-5A mice (N = 7), graphed as mean percentage of TSC2-WT littermates (N = 7) ± SEM. Quantification of immunoblots in Figure S4A. (B-D) Immunoblots and pS6/S6 ratio quantifications of lysates from skeletal muscle (quadriceps (A) (B) and gastrocnemius (C)) and liver (D) from mice subjected to a 12-h daytime fast, or similarly fasted then refed for 2 h. Graphed as mean percentage of fasted TSC2-WT littermates ± SEM; N = 3 fasted and 4 refed per genotype. (E) Echo-MRI lean mass measurements of male littermates at 3, 6, and 12 months of age, graphed as mean ± SEM; N = 12 TSC2-WT and 9 TSC2-5A. (F) Lean mass change of the measurements in (E) from 3 to 6 months of age, graphed as mean ± SEM. (G) Skeletal muscle weights in 16-week-old male TSC2-5A mice (N = 8), graphed as mean percentage of TSC2-WT littermates (N = 6) ± SEM. (H, I) Pooled (H) and binned (I) fiber cross-sectional area (CSA) distribution of the skeletal muscles of 16-week-old male TSC2-WT (N = 6) and TSC2-5A (N = 8) littermates. Data are represented as mean ± SEM. Statistical analysis by Mann-Whitney or ordinary two-way ANOVA with Tukey’s multiple comparisons tests. *p < 0.05, **p < 0.01, ***p < 0.001, ****p < 0.0001.

As mammalian lean mass is comprised of ∼40% skeletal muscle^64^, we determined whether the overall body weight changes in the TSC2-5A mice reflected a specific decrease in lean mass. Indeed, a significant decrease in lean mass was measured at 3, 6, and 12 months of age in the TSC2-5A males compared to their WT littermates (**Fig-4E**), with no significant change in fat mass (**Fig-S4B)**. The substantial lean mass gain observed between 3 and 6 months of age in TSC2-WT mice was reduced by 15% in the TSC2-5A mice (**Fig-4F**), suggesting a general decrease in muscle growth in these mice. Consistent with the delayed onset of the body weight phenotype (**Fig-3B**), female TSC2-5A mice also displayed a significant decrease in lean mass at 12 months of age, but not at 3 months (**Fig-S4C**). Like the gastrocnemius (shown in **Fig-3E**), the quadriceps and tibialis anterior were decreased in weight in the TSC2-5A mice, whereas no measurable change was observed in the small soleus muscle (**Fig-4G**). These changes in muscle weights were driven by a decrease in myofiber size, as measured by cross-sectional area, in the gastrocnemius and quadriceps muscles, but not the soleus (**Fig-4H**). A significant shift toward smaller fibers in both muscles of the TSC2-5A mice compared to TSC2-WT was observed (**Fig-4I**). Altogether, the differences observed between primary MEFs, brain, skeletal muscle, and liver demonstrate a signal- and tissue-specific dependency on TSC2 phosphorylation for full mTORC1 activation and growth.

### TSC2-5A myotubes display a cell intrinsic defect in mTORC1 signaling and protein synthesis

To determine whether the skeletal muscle and myofiber phenotypes described were associated with muscle cell-intrinsic changes, we isolated satellite cells from the muscles of TSC2-WT and 5A mice and differentiated them into myotubes *in vitro*. There was no apparent difference in myotube differentiation between the cultured TSC2-WT and −5A satellite cells (**Fig-5A**). However, insulin-stimulated mTORC1 signaling (S6K1 phosphorylation and 4EBP1 mobility shift) was severely blunted in TSC2-5A compared to -WT myotubes, despite strong activation of AKT over this time course (**Fig-5B**). As in neurons stimulated with BDNF, a small degree of mTORC1 activation with insulin was detected in TSC2-5A myotubes that was sensitive to an AKT inhibitor, but not a MEK inhibitor, suggesting a minor role for parallel AKT-dependent mechanisms for mTORC1 activation in this setting (**Fig-5C**). Previous studies found that mTORC1 is a key regulator of muscle mass downstream of AKT, at least in part by controlling protein synthesis^65^. Indeed, mirroring the effects on mTORC1 signaling, protein synthesis, as scored by relative incorporation of puromycin, was decreased in TSC2-5A myotubes both under serum starvation and insulin stimulation compared to TSC2-WT myotubes (**Fig-5D,E**). The small insulin-stimulated increase in puromycin incorporation detected in TSC2-5A myotubes was still lower than that detected in unstimulated TSC2-WT myotubes (**Fig-5D,E**). These results demonstrate that phosphorylation of TSC2 promotes muscle mTORC1 activation, protein synthesis, and growth.

**Figure 5:**
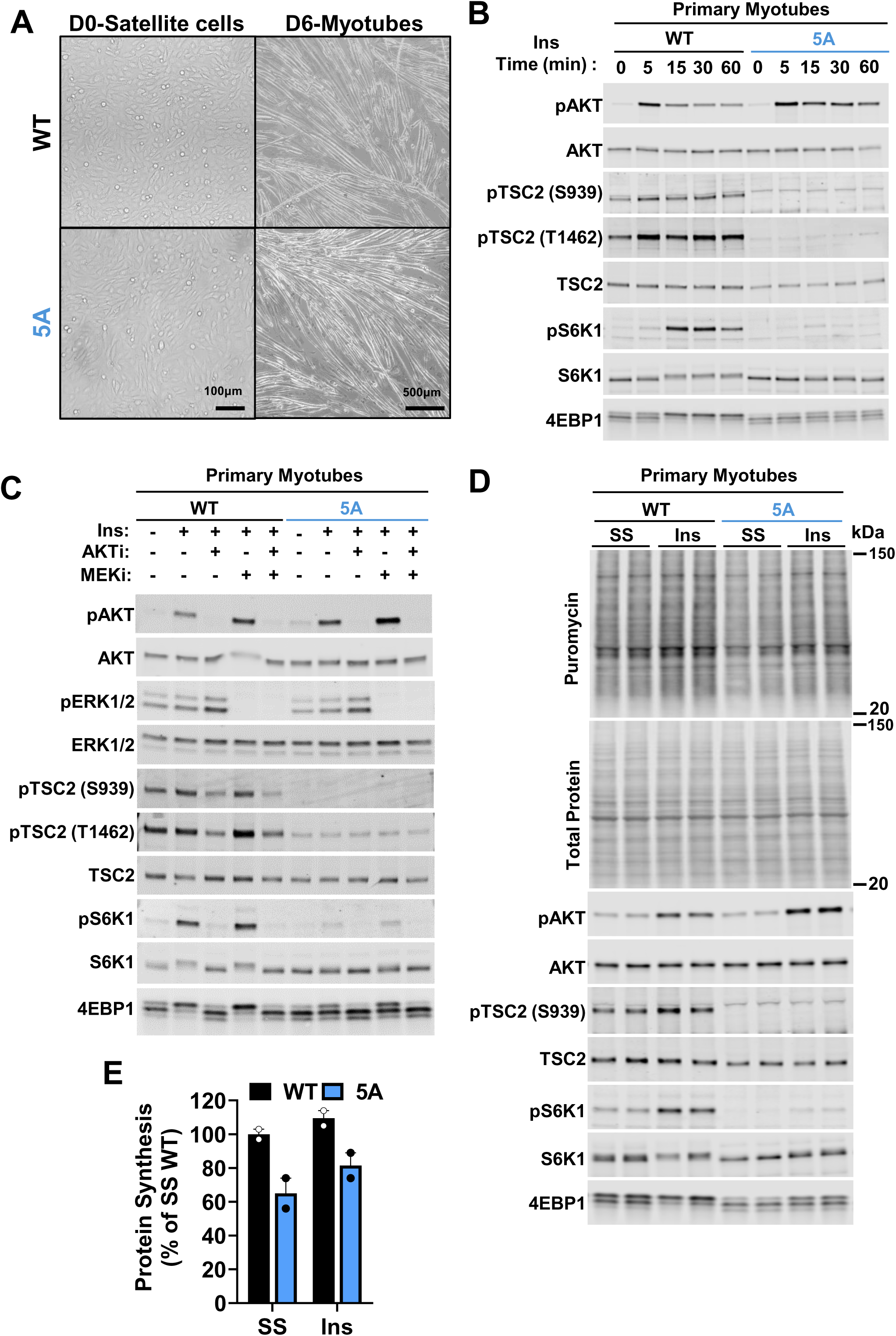
TSC2-5A myotubes display a cell intrinsic defect in mTORC1 signaling and protein synthesis. (A) Bright-field images of isolated satellite cells from TSC2-WT and TSC2-5A skeletal muscle before (D0) and after 6 days of differentiation into myotubes (D6). Scale bars: satellite cells, 100 µm; myotubes, 500 µm. (B) Representative immunoblots of lysates from primary TSC2-WT and TSC2-5A myotubes serum-starved for 6 h and then stimulated with time courses of insulin (100 nM). (C) Representative immunoblots of lysates from primary TSC2-WT and TSC2-5A myotubes serum-starved for 6 h followed by a 15-min stimulation with insulin (100 nM) after a 30-min pretreatment with vehicle (-), MK2206 (2 μM, AKTi), trametinib (5 μM, MEKi), or both, as indicated. (D) Representative immunoblots of lysates from primary TSC2-WT and TSC2-5A myotubes serum-starved for 6 hours and then stimulated with insulin (100 nM) for 1.5 h. Puromycin was added for the final 30 min of insulin stimulation to measure protein synthesis. N = 2 technical replicates. (E) Quantification of puromycin incorporation in TSC2-5A myotubes, graphed as percentage of TSC2-WT ± SEM. N = 2 independent biological experiments, each with 2 technical replicates.

## Discussion

This study describes a genetic mouse model designed to define the role of the TSC complex as a downstream target of growth-factor regulated PI3K-AKT signaling in the physiological *in vivo* regulation of mTORC1 and its effects on tissue growth. Unlike previous mouse models with complete loss of the essential components of the TSC complex, TSC1 and TSC2, which are embryonic lethal^38,41^ or, when deleted in a tissue-specific manner, give rise to constitutive maximal activation of mTORC1^22,43–48^, the TSC2-5A model represents a partial gain-of-function setting for the TSC complex in that it cannot be inhibited by the growth factor-stimulated protein kinases, such as AKT, which phosphorylate these 5 specific sites on TSC2. The TSC2-5A mice are viable, appear to develop normally, and are born at expected mendelian ratios. While this finding was unexpected, it is consistent with genetic studies in *Drosophila* lacking AKT phosphorylation sites on TSC complex components, which are also developmentally viable^58,59^. However, while TSC2-5A mice exhibit decreased postnatal body weight and organ-specific growth, the similar *Drosophila* models maintained normal body weight, wing size, and cell size, suggesting key differences in tissue growth regulation between species kept under standard laboratory conditions.

The normal embryonic development of TSC2-5A mice correlates with unaltered mTORC1 signaling in whole embryo extracts, suggesting that other mechanisms of mTORC1 regulation are dominant or redundant during mammalian embryogenesis. For instance, it is possible that other AKT-dependent mechanisms contribute, such as PRAS40 phosphorylation^12–15^. However, unlike *Tsc1* or *Tsc2* knockout, *Pras40^−/−^* mice are viable, suggesting that this mode of regulation is also dispensable for development^66^. Importantly, TSC2-WT and TSC2-5A embryos showed no difference in TSC2 phosphorylation at S664, a site attributed to ERK activity^19^, leaving open the possibility that alternative growth factor signaling pathways may contribute to mTORC1 activation during embryonic growth. It is also possible that the mechanisms of nutrient regulation of mTORC1, which occur independent of the 5 AKT phosphorylation sites on TSC2, are sufficient for the proper control of mTORC1 signaling during development.

It is interesting to consider the major growth phenotypes of the TSC2-5A mice, including decreased muscle fiber and brain size, in light of other genetic models influencing the activation state of mTORC1. For instance, whole body knockout of *Tbc1d7*, the only non-essential component of the TSC complex, leads to a specific increase in muscle fiber and brain size, associated with a modest elevation in mTORC1 signaling^42^, thus emphasizing the relative sensitivity of the growth of these tissues to changes in mTORC1 regulation. Genetic perturbations leading to stronger tissue-specific changes in mTORC1 signaling that either fully activate (*Tsc1* or *Tsc2* loss)^67,68^ or fully inhibit (*Rptor* loss)^62^ mTORC1 signaling in neurons leads to a pronounced increase (megalencephaly) or decrease (microcephaly), respectively, in brain size. The critical role of mTORC1 in promoting muscle growth and fiber size is supported by the muscle-specific knockouts of *Rptor*^53^ or *Mtor*^69^, which exhibit early-onset myopathy. Interestingly, however, deletion of either Raptor or mTOR specifically in adult muscle does not affect muscle mass in sedentary mice but leads to defects in overload-induced hypertrophy^70^. These findings suggest that mTORC1 may play different roles in developing and adult skeletal muscle.

Tissue-specific deletion of essential components of mTORC1 signaling in either the brain or skeletal muscle results in more severe phenotypes than those observed in TSC2-5A mice. This difference aligns with the fact that the TSC2-5A model disrupts a specific mode of mTORC1 activation rather than causing its complete inhibition. Indeed, a modest but significant induction of growth factor-mediated mTORC1 activation is still detectable in primary cells isolated from TSC2-5A mice, which could explain the absence of more pronounced defects. These alternate routes of mTORC1 activation in TSC2-5A cells may be due to other AKT-dependent mechanisms, such as the phosphorylation of PRAS40 or perhaps additional, uncharacterized AKT-mediated phosphorylation sites on the TSC complex. Another possible explanation includes crosstalk between the growth factor signaling and nutrient-sensing arms of mTORC1 regulation, for instance by AKT controlling nutrient uptake in some contexts^36^. Additionally, AKT has been found to phosphorylate the DEPDC5 component of GATOR1, a nutrient-sensing GAP complex for the Rag GTPases, which has been suggested to contribute to mTORC1 regulation in cells^71^. The *in vivo* characterization of alternate mechanisms of growth-factor mediated mTORC1 activation, both AKT-dependent and independent, awaits further investigation but is likely to vary between different cell and tissue types and stimuli.

Unlike skeletal muscle, liver size and feeding-induced mTORC1 signaling was found to be indistinguishable between TSC2-WT and −5A mice. These findings are aligned with a previous study showing that induction of hepatic mTORC1 signaling upon refeeding was normal in mice with liver-specific deletion of the insulin receptor^72^. Together with our results, this suggests that additional upstream signals, independent of insulin and AKT-mediated phosphorylation of TSC2, can contribute to mTORC1 activation and growth in the liver and potentially other organs not displaying weight reduction in the TSC2-5A mice. Liver size is known to be dynamically regulated by nutrient status, with liver weight decreasing by ∼25% upon fasting^52^. mTORC1 signaling likely contributes to changes in liver size as mice with constitutive hepatic mTORC1 activation through TSC1 loss have larger livers that do not shrink upon fasting, while mice with liver-specific Raptor deletion have smaller livers both in the fed and fasted state^52^. Interestingly, mouse models with either liver-specific *Tsc1* loss^45^ or active Rag GTPase signaling^73^ display sustained liver mTORC1 signaling upon fasting, indicating that the dynamic regulation of mTORC1 in the liver by fasting and feeding involves input from both the growth factor-stimulated TSC-Rheb and nutrient-sensing Rag branches. However, the dominant mechanisms contributing to hepatic mTORC1 regulation remain unclear. The heterogeneous nature of liver tissue with gradients of nutrients and hormones across liver zones^74^ might exert additional spatial cues into the regulation of hepatic mTORC1 signaling, as has been demonstrated for mTORC1 regulation by the amino acid leucine^75^.

The results described here demonstrate that AKT-mediated phosphorylation of TSC2 is a key signal promoting mTORC1 activation and the growth of some, but not all, organs. Availability of the conditional TSC2-5A mouse model will allow further investigation into the physiological and tissue-specific functions of hormone- and growth factor-induced mTORC1 signaling in a mammalian system. This is particularly interesting in the areas of metabolic homeostasis and aging, where mTORC1 is known to play key roles^76–78^,. How mTORC1 activation upon AKT-mediated TSC2 phosphorylation contributes to physiological and pathological phenotypes and which tissues are relevant can be investigated with various tissue-specific TSC2-5A models in future studies.

### Limitations of the study

While the whole-body TSC2-5A mice serve as an unbiased screen to reveal the tissues where AKT-mediated phosphorylation of TSC2 is a major mode of mTORC1 activation, the systemic nature of the model limits our interpretations, as some changes observed could stem from tissue-extrinsic factors. While the decreased brain and skeletal muscle weight characterized in the TSC2-5A mice is correlated with cell-intrinsic defects in isolated neurons and myotubes, it remains unclear whether the inability to phosphorylate these 5 sites on TSC2 in other cell types or tissues is directly contributing to these or other growth phenotypes. Unlike other tissues that are distinct between the TSC2-WT and −5A mice, we see an increase in the weight of specific fat pads from the TSC2-5A mice without an overall significant change in fat mass. While we do not currently know the underlying cause of this difference, it seems likely that this specific increase is secondary to a change elsewhere in these animals. Furthermore, TSC2-5A males display an earlier onset and more pronounced body-weight phenotype than females from the same cohort. The basis for this sex-specific difference in body weight with age observed in the TSC2-5A mice remains unknown. Future studies will help address these limitations and take advantage of the conditional nature of these mice to generate tissue-specific TSC2-5A models that facilitate a deeper investigation of the *in vivo* role of AKT-mediated phosphorylation of TSC2 for the control of mTORC1 and for the characterization of tissue-specific downstream functions.

## Acknowledgments

We thank Drs. Eva Tsaousidou and Karen Inouye for advice and technical assistance and our colleagues at the Harvard Genome Modification Facility. This study was supported by a Glenn Foundation for Medical Research Postdoctoral Fellowship (Y.C.), National Institutes of Health grants T32-DK128781 (S.C.L., K.C.K), F31-DK128873 (K.C.K), R35-CA197459 (B.D.M.), and P01-CA120964 (B.D.M.), and Damon Runyon Cancer Research Foundation Merck Fellowship DRG-#2443-21 (M.Y.C.).

## Author contributions

Conceptualization, Y.C., S.M. and B.D.M.; investigation, Y.C., S.C.L., K.C.K., M.Y.C., J.S., B.S., J.P., S.C., A.D.A., E.R.M, C.L.S., S.M., and V.B.; writing, Y.C., K.C.K., and B.D.M.; funding acquisition, Y.C., K.C.K., M.Y.C., and B.D.M.; resources, B.D.M. and M.S.; supervision, B.D.M., and M.S.

## Declaration of interests

M.S. reports grant support from Biogen, Astellas, Bridgebio, and Aucta, and he has served on Scientific Advisory Boards (SAB) for Roche, SpringWorks Therapeutics, and Alkermes and is currently on the SABs for Neurogene, Jaguar Gene Therapy, and Noema. All other authors declare no competing financial interests.

## Methods

### Mice and diets

All animal studies were reviewed and approved by the Harvard Medical Area Standing Committee on Animals IACUC, an AAALAC International and USDA-accredited facility. All animal care and protocols were in accordance with the principles of animal care and experimentation in the Guide for the Care and Use of Laboratory Animals. Mice were separated by gender, group housed (3-4 per cage) in temperature controlled, pathogen-free facilities with a 12:12 h light:dark cycle in standard static microisolator top cages. Autoclaved food (Lab Diet #5053, St Louis MO) and water were provided ad libitum

### Generation of the Rosa26-LSL-Flag-TSC2-WT and -TSC2-5A transgene knock in mice

The full length human Flag-TSC2-WT or Flag-TSC2-5A (S939A, S981A, S1130A, S1132A, T1462A) cDNAs, derived from TSC2 transcript P49815-4 and previously described^79^, were recombined into intron 1 of the mouse *Rosa26* locus using the targeting vector STOP-eGFP-ROSA26TV (Addgene #11739)^56^. The TSC2-WT and −5A targeting vectors were introduced into C57BL/6-129 Hybrid ES cells and G418-resistant ES clones were screened for homologous recombination with the *Rosa26* locus by PCR using the following primers (see **Fig-S1D** for primer position): 5′ junction forward P1 (5’-ttggtgcgtttgcggggatg-3’), reverse P2 (5’-catcaaggaaaccctggactactg-3’); 3′ junction forward P3 (5′-atggccacaaccatggtgagc-3’), reverse P4 (5’-actcgcgacactgtaatttcatac-3’). Chimeric mice were generated by injection of one ES cell clone of each genotype into C57BL/6J blastocysts and transferred into recipient foster mothers. Chimeric mice were mated with C57BL/6J mice to obtain germline transmission. Genotyping for the presence of the transgene (Tg) or wild type *Rosa26* (R26) locus was performed using the following 3 primers: P5 (5’-ttgctctcccaaagtcgctctga-3’), P6 (5’-aatctgtgggaagtcttgtcc-3’) and P2 (5’-catcaaggaaaccctggactactg-3’). To confirm the presence or Cre-mediated deletion of the LSL cassette the following primers were used: Presence: P7 (5’-gtccagggtttccttgatgatgt-3’) P8 (5’-ttcctcgtgctttacggtatcgc-3’), deletion: P7 and P9 (5’-cacattccatgctcagttctctca-3’). As the S939A mutation in the TSC2-5A transgene was designed to eliminate an SpeI restriction enzyme site, we took advantage of this mutation to differentiate between the TSC2-WT and −5A transgenes between littermates. The following primers were used to amplify the S939 region: P10 (5’-cgtcatagccatgtggttca-3’), P11 (5’-atcatgtccagacaggtttcc-3’); the PCR product was then purified (Qiagen, 28106), digested with SpeI enzyme (NEB, R3133L), and resolved on an agarose gel. Location of the primers and representative PCR results are displayed in **Fig-S1D-G**.

### Isolation and culturing of primary mouse embryonic fibroblasts

The following protocol was adapted from Durkin et al.^80^. Briefly, E13.5–15.5 embryos were harvested and transferred individually to Petri dishes containing sterile phosphate-buffered saline (PBS). Each embryo was kept separate to generate independent MEF lines. The head, heart, and liver were removed, and the remaining embryo was placed into a Petri dish with 0.25% trypsin-EDTA. The embryos were then minced into 1–2 mm pieces using a razor blade and placed in a 37 °C tissue culture incubator for 10 min, followed by pipetting up and down several times with a 5 ml pipette before an additional 5–10 min incubation at 37 °C. The dispersed cell suspension was then transferred to a 50 ml conical tube containing 20-ml warmed Dulbecco’s Modified Eagle Medium (DMEM 4.5 g/L glucose, without sodium pyruvate; Corning, 15-017-CV), 10% FBS, 1X Pen/Strep, and 2 mM glutamine. The cell suspension was allowed to sit for 5 minutes to let larger fragments settle, and the supernatant was transferred to a 60-mm cell culture dish. The following day, fresh medium was added, and the cells were monitored for confluency. When confluent, the cells were transferred to 10 or 15 cm dishes to expand. Cells of fewer than 7 passages were used for the described experiments. For signaling experiments, primary MEFs were serum-starved for 16 h, followed by stimulation with FBS (10%), EGF (100 ng/ml), or insulin (100 nM) after a 30-minute pretreatment with vehicle, MK2206 (2 μM), trametinib (5 μM), both, or rapamycin (20 nM), where indicated.

### Primary neuron isolation and culture

The cortex and hippocampus were dissected from the brains of E16.5 TSC2-WT and −5A mouse embryos and dissociated in 0.25% trypsin (Thermo #25200114) for 15-min at 37C. Four washing steps were performed, two in Hank’s balanced salt solution buffer (Life Tech #14170-120), one in Neurobasal A medium (Life Tech #10888-022) supplemented with 1% Glutamine, 2% B27 (Invitrogen #17504-044), 1% penicillin/streptomycin and 100 mM b-mercaptoethanol (growth media), and one in Neurobasal A supplemented with 10% horse serum (Sigma #H1138). Tissue pieces were then triturated in growth medium using sterile fire-polished Pasteur pipettes (VWR #14672-380). Single-cell suspensions of hippocampal neurons were plated on glass coverslips previously washed with 100% methanol, 70% ethanol and 100% ethanol, dried, and coated with 30 ng/mL poly-ornithine (Sigma #P3655). For biochemical experiments, cortical neurons were plated in 12-well plates at a density of 300,000 cells per well. For microscopy experiments, hippocampal neurons were plated in 24-well plates at 40,000 cells per well. Neurons were maintained in a humidified incubator at 37 °C and 5% CO2 for 4 to 7 Days In Vitro (DIV) before performing signaling and immunofluorescence assays. For signaling experiments, primary neurons were starved in HBSS media for 4 h followed by BDNF (100 ng/ml) or IGF1 (100 ng/ml) stimulation with a 30-min vehicle, MK2206 (2 μm), trametinib (5μm), both, or rapamycin (20 nM) pretreatment where indicated.

### Primary hepatocyte isolation and culture

Primary hepatocytes were isolated from C57/BL6J male mice at 8 weeks of age (Jackson Labs) as previously described^81^. Portal vein perfusion of buffer A (10 mM HEPES, 150 mM NaCl, 5 mM KCl, 5 mM glucose, and 2.5 mM sodium bicarbonate, and 0.5 mM EDTA, pH 8.0) was performed for 5–7 min at a rate of 5 ml/min followed by buffer B (buffer A minus EDTA, with 35 mM CaCl2 and Liberase TM, Sigma–Aldrich, 5401127001) for 5–7 min at a rate of 5 ml/min. Livers were placed in Dulbecco’s Modified Eagle Medium (DMEM 4.5 g/L glucose, w/o sodium pyruvate; Corning, 15-017-CV), 2.5% FBS, 1X Pen/Strep, and Glutamax (ThermoFisher, 35050061) or Glutagro (Corning, 25-015-CI), and the liver capsule was disrupted to release hepatocytes. Hepatocytes were centrifuged at 1000 rpm for 5 min and then resuspended in a mixture of 10 ml DMEM, 9 ml Percoll (Sigma–Aldrich, P4937), and 1 ml 10x PBS. Following centrifugation at 1000 rpm for 7 min, hepatocytes were washed once with media, centrifuged at 1000 rpm for 5 min, and then resuspended in 10 ml media per liver. Viability was determined by Trypan blue exclusion and cell counts. Hepatocytes were plated at 1.25–1.5 × 10^6^ cells/well in 6-well collagen-coated dishes (BioCoat, Corning). After 4–6 h, media were changed to serum-free media (DMEM, 2 mM l-glutamine) overnight followed by insulin stimulation (100 nM human insulin, Sigma–Aldrich, I9278), with a 30-min vehicle, MK2206 (2 μm), trametinib (5 μm), both, or rapamycin (20 nM) pretreatment, where indicated.

### Satellite cell isolation, culture, and differentiation into myotubes

The following protocol was adapted from Wright et al.^82^ In brief, lower limb muscles were dissected from newly sacrificed 8-week-old male mice, finely minced, and digested in DMEM (4.5 g/L glucose) containing 2mg/mL collagenase type II (ThermoFischer Scientific; 17101015) and 0.5 mg/mL dispase (ThermoFischer Scientific; 17105041) for 60 min at 37°C. During digestion, solutions were pipetted periodically in descending order using 25 mL, 10 mL, 5 mL, and 2 mL serological pipettes until the solution passed through without resistance. Digestion was halted using 2X solution volume of growth medium (DMEM [4.5 g/L glucose]/Ham’s F-10 supplemented with 20% FBS, 1Xpenicillin/streptomycin [R&D Systems; B21210]) then passed through a 100-μm filter. Samples were centrifuged at 1500 rpm for 10 min. Red blood cells were lysed for 2 min at room temperature using Red Blood Cell Lysis Solution (Milteny Biotec; 130-094-183). Samples were passed through a 40-μm filter, then centrifuged as above. Samples were subjected to Debris Removal Solution (Milteny Biotec; 130-109-398) according to manufacturer instructions. Samples were depleted of non-satellite cell populations using MACS Mouse Satellite Cell Isolation Kit (Milteny Biotec; 130-104-268) according to manufacturer instructions. Satellite cells were plated on laminin-511 (Biogems; RL511) coated 10 cm plates in growth medium supplemented with 10 ng/mL bFGF (PeproTech; 450-33), refreshed daily. Satellite cells were expanded in growth medium and passaged prior to any region reaching greater than 60% confluence. All experiments were performed at or before passage 4. Cells were plated on laminin-511 (Biogems; RL511) coated 6-well plates at 200,000 cells per well and allowed to recover for one day in growth medium. 24 h after plating, media was changed to differentiation medium (DMEM [1 g/L glucose]/Ham’s F-10 supplemented with 2% FBS and 1X penicillin/streptomycin [R&D Systems; B21210]), changed daily for 5 days. After 5 days, medium was changed to experimental medium (Ham’s F-12 supplemented with 4% HS and 1X penicillin/streptomycin [R&D Systems; B21210]) for 24 hours, after which experiments were initiated. DIC images of differentiated myotubes were taken on a tabletop light microscope. For insulin signaling experiments, myotubes were serum starved in Ham’s F-12 supplemented with 1X penicillin/streptomycin [R&D Systems; B21210] for 6 h. Following serum starvation, cells were treated with 100nM insulin for 15 minutes. 30 minutes prior to insulin treatment, cells were treated with or without the following MK2206 (2 μm), trametinib (5 μm), both, or rapamycin (20 nM), where indicated.

### Confocal immunofluorescence microscopy of primary neurons

Cultured primary hippocampal neurons were fixed on glass coverslips in warm 4% paraformaldehyde (PFA) in PBS for 15 min at room temperature and permeabilized and blocked in Intercept PBS blocking buffer (Licor #927-70001) with 0.1% Triton for 1 h at room temperature. Coverslips were incubated in blocking buffer containing primary antibodies 1 h at room temperature. Following three 5-min washes with PBS, coverslips were incubated in blocking buffer with secondary antibodies for 30 min at room temperature. Following three 10-min washes with PBS, with DAPI (Thermo #62248) added to the last wash, coverslips were mounted to slides using Fluoromount-G (Southern Biotech #0100-01). antibodies to MAP2 (Synaptic Systems #188004) and guinea pig IgG (Cy3-conjugated; 1:500; Jackson ImmunoResearch #706-165-148) were used as primary and secondary antibodies, respectively. Slides were imaged with a Nikon Ti-E inverted microscope (Nikon Instruments, Melville, NY), using a 60x objective lens with a Zyla 4.2 cMOS camera and NIS elements software for acquisition parameters, shutters, filter positions and focus control. Neuronal soma area was determined by manually measuring MAP2 staining in Fiji.

### Confocal immunofluorescence microscopy of skeletal muscle

The following was adapted from Wu et al.^83^. Skeletal muscles were dissected and immediately frozen in isopentane cooled in liquid nitrogen for 10-30 sec depending on muscle size. Samples were immediately placed on dry ice then transferred to −80°C for storage. Frozen tissues were transferred on dry ice from −80°C to −20°C cryostat (Microtome HM 505N Cryostat). After tissue temperature equilibrated to cryostat temperature, they were cut transversely at the mid-belly. Half of the tissue was embedded in Tissue-Tek O.C.T. Compound (Sakura; 4583), and the other was returned to −80°C. Serial sections were collected at 10 μm thickness onto Superfrost microscope slides (VWR; 48311-703) and stored at −80°C until staining.

The following was adapted from Esper et al.^84^. To measure cross-sectional area (CSA) of the gastrocnemius, quadriceps, and soleus muscles, slides were removed from −80°C and allowed to dry for 30 min at RT. Slides were then fixed in 4% paraformaldehyde for 15 min at 4°C. Slides were washed 3 times for 5 min each in 1X PBS. Heat-induced antigen retrieval (HIAR) was performed in 1-L 10 mM citrate buffer (2.94g sodium citrate tribasic, dihydrate, 0.5mL Tween 20, pH 7, to 1L in water). Slides were place in pre-warmed citrate buffer and microwaved to maintain a low boil for 10 min. Citrate buffer was washed out with cold, running, distilled water for 10 min. Slides were placed in permeabilization buffer (1X PBS, 0.1 M glycine, 0.1% Triton-X, syringe filtered) for 10 min, then blocking buffer (1X PBS, 5% goat serum (Jackson ImmunoResearch; 005-000-121), 2% bovine serum albumin (Sigma-Aldrich; A9647), 1:40 Mouse on Mouse (Vector Laboratories; MKB-2213-1), syringe filtered) for 1 h at room temperature. HIAR was performed for quadriceps and gastrocnemius muscles only; soleus sections proceeded directly from fixation to permeabilization. Slides were incubated overnight at 4°C with Alexa Fluor 594-conjugated wheat germ agglutinin (WGA, ThermoFischer; W11262), Type 2B fibers were stained in the gastrocnemius and quadriceps with BF-F3 (concentrate, Developmental Studies Hybridoma Bank; 1:40). Slides were washed 3 times for 5 min each in 1X PBS, then incubated for 60 min at room temperature with Alexa Fluor 488 Goat anti-Mouse IgM, μ-chain specific (Jackson ImmunoResearch; 115-545-075). Slides were washed 3 times for 5 each with 1X PBS, then mounted in VectaShield Hardset with DAPI (Vector Laboratories; H-1500-10). Slides were incubated at room temperature for 30 min, then sealed with clear nail polish.

Slides were imaged on Nikon Eclipse Ti microscope with X-Cite 120LED lamphouse. Images were captured on an ANDOR Zyla 5.5 SCMOS camera using NIS-Elements software. All images were captured under 20X magnification under 27% iris intensity. Images for gastrocnemius and quadriceps were composed of 3×3 fields (whole soleus was imaged with 4×4 field) with 15% overlap and stitched on NIS-Elements software. Three composite images were taken of the gastrocnemius, one within each head and one deep image. Four composite images were taken of the quadriceps, one image composed primarily of each individual muscle (vastus lateralis, vastus intermedius, vastus medialis, and rectus femoris). Type 2B staining and tissue landmarks were used to ensure images came from the same regions of the gastrocnemius or quadriceps. One composite image of the entire soleus was captured and analyzed.

All images used for cross sectional area (CSA) calculations were analyzed in a blinded fashion, using solely the TRITC channel, WGA images. Brightness of images were uniformly adjusted for analysis. Semi-automatic muscle analysis using segmentation of histology (SMASH) was used to analyze CSA of each image. Images were analyzed with the following parameters, pixel size (μm/pixel) 0.326, minimum fiber area (μm^2^) 400, maximum fiber area (μm^2^) 9000, maximum eccentricity 0.95, minimum convexity 0.8. Manual outlining was used to compensate for differences in WGA intensity. Myofibers damaged by sectioning, overlapping myofibers, incomplete myofibers, and any non-myofiber selections were manually removed as necessary.

### Protein Extraction and Immunoblotting

Protein extracts were prepared from tissues and cells using ice-cold NP-40 lysis buffer (20 mM Tris pH8, 150 mM NaCl, 10% Glycerol, 1% NP-40 (Igepal CA-630), protease (Sigma #P8340) and phosphatase (Sigma #P2850) inhibitors), Samples were centrifuged at 20,000 g for 15-min at 4 °C, and protein concentration in the supernatant was determined by Bradford assay (Thermo Fisher Scientific). Equal amounts of protein (Cells: 5-10 μg, Tissues: 30-40 μg) were loaded per well and separated by SDS-PAGE, transferred to nitrocellulose membranes, and immunoblotted with indicated primary antibodies. Primary antibodies: phospho (P)–S6K1 T389 (Cell Signaling Technologies (CST), 9234), S6K1 (CST, 2708) P-AKT S473 (CST, 4060), Pan-AKT (CST, 4691), P–S6 S240/44 (CST, 2215), S6 (CST, 2217), 4E-BP1 (CST, 9644), TSC2 (CST, 3612), P-TSC2 S664 (CST, 40729), P-TSC2 S939 (CST, 3615)), P-TSC2 S1387 (CST, 9644), P-TSC2 T1462 (CST, 3617), ERK1/2 (CST, 9102) P-ERK1/2 (CST, 4370), PRAS40 (CST, 2610), P-PRAS40 (CST, 2997), mTOR (CST, 2983), P-mTOR S2448 (CST, 2971), AMPKα (CST, 2603), P-AMPKα (CST, 2535)and Puromycin (Millipore, MABE343). Secondary antibodies were IRDye 800CW donkey anti-rabbit IgG (LI-COR, 925-32213) and IRDye 800CW donkey anti-mouse IgG (LI-COR, 926-32212). Immunoblots were imaged with a LI-COR Odyssey CLx imaging system (LI-COR Biosciences) and quantified using the LI-COR software.

### Protein synthesis assay

The following was adapted from Schmidt et al.^85^. Myotubes were serum starved in Ham’s F-12 supplemented with 1X penicillin/streptomycin [R&D Systems; B21210] for 6 h. Following serum starvation, cells were treated with 100 nM insulin for a total of 1.5 h with addition of 1 μM puromycin (Sigma, P7255) for the final 30 min. Protein was extracted and immunoblotted as described above. Total protein was assessed using Revert™ 700 Total Protein Stain (LI-COR, 926-11011) in the 700 nm channel using LI-COR Odyssey CLx imaging system (LI-COR Biosciences). Puromycin incorporation was assessed by immunoblotting for puromycin (Millipore, MABE343) on the same membrane, assessed in the 800 nm channel. Protein synthesis was quantified using ImageJ by normalizing puromycin intensity to total protein intensity.

### Gene expression analysis

RNA was isolated from whole E15.5 embryos using TRIzol Reagent (ThermoFisher 15596026). 1 μg RNA was reverse transcribed using the Advanced cDNA Synthesis Kit (Bio-Rad). Skirted plates and iTaq SYBR green for qPCR were purchased from Bio-Rad. The QPCR analysis was performed in biological duplicates or triplicates and with technical duplicates. Analysis was performed using the CFX96 Real-Time PCR Detection System. Samples were normalized to Rplp0 (36b4) for ΔΔCt analysis using the Bio-Rad CFX96 software. Primers: human-TSC2-forward (5’-GGCTCTATTCTCTCAGGAACTC-3’), human-TSC2-reverse (5’-GATGGACAGGACGATCTCATAG-3’), mouse-TSC2-forward (5’-GCATGCAGTTCTCACCTTAT-3’), mouse-TSC2-reverse (5’-GGGTAGTCCTTGATCACTTTG-3’), mouse-36b4-forward (5’-AGATGCAGCAGATCCGCAT-3’), mouse-36b4-reverse (5’-GTTCTTGCCCATCAGCACC-3’).

### TSC1 immunoprecipitation assay

Protein extracts were prepared from whole E15.5 embryos using 1% Triton X-100 lysis buffer (40mM HEPES pH 7.4, 120 mM NaCl, 1 mM EDTA, 1% Triton X-100, 1X protease inhibitor cocktail (Sigma), 1X phosphatase inhibitor cocktail (Roche)). Samples of 500 μg protein extract were pre-cleared with 50 μl of 50% protein A/G beads (Pierce) for 30 min on a rotator at 4 °C and then transferred to a new tube with 5 μl TSC1 antibody (R&D Systems, MAB4379) or 25 μl mouse IgG Control (SC-2025) and incubated overnight on a rotator at 4 °C. 50 μL of 50% pre-washed protein A/G beads (Pierce) were added to each sample and incubated for 1 h on a rotator at 4 °C. The beads were then washed 5 times for 5 min each with lysis buffer (0.1X protease/phosphatase inhibitors) on a rotator at 4 °C, followed by final centrifugation. Pelleted immune complexes on beads were resuspended in 2X Laemmli sample buffer and denatured at 95 °C for 5 min. Samples were then subjected to SDS-PAGE and immunoblotting, where 10 μg of protein extract served as the input controls and 40% of each immunoprecipitation was loaded.

### EchoMRI scan

Animals were placed in a ventilated clear plastic holder (vertical bore) without sedation or anesthesia with restricted movement during the procedure. The holder is inserted into a tubular space within the EchoMRI™ system to measure lean and fat mass. After scanning, animals were immediately returned to normal caging.

### Statistical analyses

Statistical analyses were performed using GraphPad Prism 10 software. Data are presented as mean ± standard error of mean (SEM). When analyzing two groups, a Mann-Whitney test was used. For comparison between signaling time courses, a repeated measurement two-way ANOVA with Greenhouse–Geisser correction was performed. For body weight measurements over time, a mixed model effect with Šídák’s multiple comparisons test was performed. For comparisons between three or more groups, a Kruskal-Wallis or an ordinary two-way ANOVA with Tukey’s multiple comparisons test, with a single pooled variance were performed.

## References

1. James, D.E., Stöckli, J., and Birnbaum, M.J. (2021). The aetiology and molecular landscape of insulin resistance. Nat. Rev. Mol. Cell Biol. 2021 2211 22, 751–771. 10.1038/s41580-021-00390-6.

2. Saltiel, A.R. (2021). Insulin signaling in health and disease. J. Clin. Invest. 131. 10.1172/JCI142241.

3. Valvezan, A.J., and Manning, B.D. (2019). Molecular logic of mTORC1 signalling as a metabolic rheostat. Nat. Metab. 2019 13 1, 321–333. 10.1038/s42255-019-0038-7.

4. Reynolds IV, T.H., Bodine, S.C., and Lawrence, J.C. (2002). Control of Ser2448 phosphorylation in the mammalian target of rapamycin by insulin and skeletal muscle load. J. Biol. Chem. 277, 17657–17662. 10.1074/jbc.M201142200.

5. Navé, B.T., Ouwens, D.M., Withers, D.J., Alessi, D.R., and Shepherd, P.R. (1999). Mammalian target of rapamycin is a direct target for protein kinase B: identification of a convergence point for opposing effects of insulin and amino-acid deficiency on protein translation. Biochem. J. 344, 427. 10.1042/0264-6021:3440427.

6. Sekulić, A., Hudson, C.C., Homme, J.L., Yin, P., Otterness, D.M., Karnitz, L.M., and Abraham, R.T. (2000). A direct linkage between the phosphoinositide 3-kinase-AKT signaling pathway and the mammalian target of rapamycin in mitogen-stimulated and transformed cells. Cancer Res. 60, 3504–3513.

7. Rosner, M., and Hengstschläger, M. (2008). Cytoplasmic and nuclear distribution of the protein complexes mTORC1 and mTORC2: rapamycin triggers dephosphorylation and delocalization of the mTORC2 components rictor and sin1. Hum. Mol. Genet. 17, 2934– 2948. 10.1093/HMG/DDN192.

8. Holz, M.K., and Blenis, J. (2005). Identification of S6 kinase 1 as a novel mammalian target of rapamycin (mTOR)-phosphorylating kinase. J. Biol. Chem. 280, 26089–26093. 10.1074/jbc.M504045200.

9. Chiang, G.G., and Abraham, R.T. (2005). Phosphorylation of mammalian target of rapamycin (mTOR) at Ser-2448 is mediated by p70S6 kinase. J. Biol. Chem. 280, 25485– 25490. 10.1074/jbc.M501707200.

10. Kovacina, K.S., Park, G.Y., Bae, S.S., Guzzetta, A.W., Schaefer, E., Birnbaum, M.J., and Roth, R.A. (2003). Identification of a proline-rich Akt substrate as a 14-3-3 binding partner. J. Biol. Chem. 278, 10189–10194. 10.1074/jbc.M210837200.

11. Oshiro, N., Takahashi, R., Yoshino, K.I., Tanimura, K., Nakashima, A., Eguchi, S., Miyamoto, T., Hara, K., Takehana, K., Avruch, J., et al. (2007). The proline-rich Akt substrate of 40 kDa (PRAS40) is a physiological substrate of mammalian target of rapamycin complex 1. J. Biol. Chem. 282, 20329–20339. 10.1074/jbc.M702636200.

12. Fonseca, B.D., Smith, E.M., Lee, V.H.Y., MacKintosh, C., and Proud, C.G. (2007). PRAS40 is a target for mammalian target of rapamycin complex 1 and is required for signaling downstream of this complex. J. Biol. Chem. 282, 24514–24524. 10.1074/jbc.M704406200.

13. Sancak, Y., Thoreen, C.C., Peterson, T.R., Lindquist, R.A., Kang, S.A., Spooner, E., Carr, S.A., and Sabatini, D.M. (2007). PRAS40 Is an Insulin-Regulated Inhibitor of the mTORC1 Protein Kinase. Mol. Cell 25, 903–915. 10.1016/J.MOLCEL.2007.03.003.

14. Haar, E. Vander, Lee, S. il, Bandhakavi, S., Griffin, T.J., and Kim, D.H. (2007). Insulin signalling to mTOR mediated by the Akt/PKB substrate PRAS40. Nat. Cell Biol. 2007 93 9, 316–323. 10.1038/ncb1547.

15. Wang, L., Harris, T.E., Roth, R.A., and Lawrence, J.C. (2007). PRAS40 regulates mTORC1 kinase activity by functioning as a direct inhibitor of substrate binding. J. Biol. Chem. 282, 20036–20044. 10.1074/jbc.M702376200.

16. Manning, B.D., Tee, A.R., Logsdon, M.N., Blenis, J., and Cantley, L.C. (2002). Identification of the tuberous sclerosis complex-2 tumor suppressor gene product tuberin as a target of the phosphoinositide 3-kinase/Akt pathway. Mol. Cell 10, 151–162. 10.1016/S1097-2765(02)00568-3.

17. Inoki, K., Li, Y., Zhu, T., Wu, J., and Guan, K.L. (2002). TSC2 is phosphorylated and inhibited by Akt and suppresses mTOR signalling. Nat. Cell Biol. 4, 648–657. 10.1038/ncb839.

18. Menon, S., Dibble, C.C., Talbott, G., Hoxhaj, G., Valvezan, A.J., Takahashi, H., Cantley, L.C., and Manning, B.D. (2014). Spatial control of the TSC complex integrates insulin and nutrient regulation of mTORC1 at the lysosome. Cell 156, 771–785. 10.1016/j.cell.2013.11.049.

19. Ma, L., Chen, Z., Erdjument-Bromage, H., Tempst, P., and Pandolfi, P.P. (2005). Phosphorylation and functional inactivation of TSC2 by Erk: Implications for tuberous sclerosis and cancer pathogenesis. Cell 121, 179–193. 10.1016/j.cell.2005.02.031.

20. Roux, P.P., Ballif, B.A., Anjum, R., Gygi, S.P., and Blenis, J. (2004). Tumor-promoting phorbol esters and activated Ras inactivate the tuberous sclerosis tumor suppressor complex via p90 ribosomal S6 kinase. Proc. Natl. Acad. Sci. U. S. A. 101, 13489–13494. 10.1073/PNAS.0405659101/ASSET/6FD259C3-D3B9-40B1-B9B2-984DDB95C335/ASSETS/GRAPHIC/ZPQ0380460150006.JPEG.

21. Dibble, C.C., Elis, W., Menon, S., Qin, W., Klekota, J., Asara, J.M., Finan, P.M., Kwiatkowski, D.J., Murphy, L.O., and Manning, B.D. (2012). TBC1D7 Is a Third Subunit of the TSC1-TSC2 Complex Upstream of mTORC1. Mol. Cell 47, 535–546. 10.1016/j.molcel.2012.06.009.

22. Kwiatkowski, D.J., Zhang, H., Bandura, J.L., Heiberger, K.M., Glogauer, M., el-Hashemite, N., and Onda, H. (2002). A mouse model of TSC1 reveals sex-dependent lethality from liver hemangiomas, and up-regulation of p70S6 kinase activity in Tsc1 null cells. Hum. Mol. Genet. 11, 525–534. 10.1093/HMG/11.5.525.

23. Jaeschke, A., Hartkamp, J., Saitoh, M., Roworth, W., Nobukuni, T., Hodges, A., Sampson, J., Thomas, G., and Lamb, R. (2002). Tuberous sclerosis complex tumor suppressor–mediated S6 kinase inhibition by phosphatidylinositide-3-OH kinase is mTOR independent. J. Cell Biol. 159, 217–224. 10.1083/JCB.JCB.200206108.

24. Goncharova, E.A., Goncharov, D.A., Eszterhas, A., Hunter, D.S., Glassberg, M.K., Yeung, R.S., Walker, C.L., Noonan, D., Kwiatkowski, D.J., Chou, M.M., et al. (2002). Tuberin regulates p70 S6 kinase activation and ribosomal protein S6 phosphorylation: A role for the TSC2 tumor suppressor gene in pulmonary lymphangioleiomyomatosis (LAM). J. Biol. Chem. 277, 30958–30967. 10.1074/JBC.M202678200/ASSET/F68BB189-D909-4957-950D-508ABBF864CA/MAIN.ASSETS/GR11.JPG.

25. Dibble, C.C., and Manning, B.D. (2013). Signal integration by mTORC1 coordinates nutrient input with biosynthetic output. Nat. Cell Biol. 15, 555–564. 10.1038/ncb2763.

26. Inoki, K., Zhu, T., and Guan, K.-L. (2003). TSC2 Mediates Cellular Energy Response to Control Cell Growth and Survival. Cell 115, 577–590. 10.1016/S0092-8674(03)00929-2.

27. Inoki, K., Ouyang, H., Zhu, T., Lindvall, C., Wang, Y., Zhang, X., Yang, Q., Bennett, C., Harada, Y., Stankunas, K., et al. (2006). TSC2 Integrates Wnt and Energy Signals via a Coordinated Phosphorylation by AMPK and GSK3 to Regulate Cell Growth. Cell 126, 955–968. 10.1016/j.cell.2006.06.055.

28. Gwinn, D.M., Shackelford, D.B., Egan, D.F., Mihaylova, M.M., Mery, A., Vasquez, D.S., Turk, B.E., and Shaw, R.J. (2008). AMPK Phosphorylation of Raptor Mediates a Metabolic Checkpoint. Mol. Cell 30, 214–226. 10.1016/j.molcel.2008.03.003.

29. Horman, S., Vertommen, D., Heath, R., Neumann, D., Mouton, V., Woods, A., Schlattner, U., Wallimann, T., Carling, D., Hue, L., et al. (2006). Insulin Antagonizes Ischemia-induced Thr172 Phosphorylation of AMP-activated Protein Kinase α-Subunits in Heart via Hierarchical Phosphorylation of Ser485/491. J. Biol. Chem. 281, 5335–5340. 10.1074/JBC.M506850200.

30. Kim, E., Goraksha-Hicks, P., Li, L., Neufeld, T.P., and Guan, K.L. (2008). Regulation of TORC1 by Rag GTPases in nutrient response. Nat. Cell Biol. 2008 108 10, 935–945. 10.1038/ncb1753.

31. Sancak, Y., Peterson, T.R., Shaul, Y.D., Lindquist, R.A., Thoreen, C.C., Bar-Peled, L., and Sabatini, D.M. (2008). The rag GTPases bind raptor and mediate amino acid signaling to mTORC1. Science (80-.). 320, 1496–1501. 10.1126/science.1157535.

32. Sancak, Y., Bar-Peled, L., Zoncu, R., Markhard, A.L., Nada, S., and Sabatini, D.M. (2010). Ragulator-rag complex targets mTORC1 to the lysosomal surface and is necessary for its activation by amino acids. Cell 141, 290–303. 10.1016/j.cell.2010.02.024.

33. Zoncu, R., Bar-Peled, L., Efeyan, A., Wang, S., Sancak, Y., and Sabatini, D.M. (2011). mTORC1 senses lysosomal amino acids through an inside-out mechanism that requires the vacuolar H+-ATPase. Science (80-.). 334, 678–683. 10.1126/SCIENCE.1207056/SUPPL_FILE/ZONCU.SOM.PDF.

34. Bar-Peled, L., Chantranupong, L., Cherniack, A.D., Chen, W.W., Ottina, K.A., Grabiner, B.C., Spear, E.D., Carter, S.L., Meyerson, M., and Sabatini, D.M. (2013). A tumor suppressor complex with GAP activity for the Rag GTPases that signal amino acid sufficiency to mTORC1. Science (80-.). 340, 1100–1106. 10.1126/SCIENCE.1232044/SUPPL_FILE/1232044BAR-PELED.SM.PDF.

35. Efeyan, A., Zoncu, R., Chang, S., Gumper, I., Snitkin, H., Wolfson, R.L., Kirak, O., Sabatini, D.D., and Sabatini, D.M. (2012). Regulation of mTORC1 by the Rag GTPases is necessary for neonatal autophagy and survival. Nat. 2012 4937434 493, 679–683. 10.1038/nature11745.

36. Manning, B.D., and Toker, A. (2017). AKT/PKB Signaling: Navigating the Network, 10.1016/j.cell.2017.04.001 https://doi.org/10.1016/j.cell.2017.04.001.

37. Onda, H., Lueck, A., Marks, P.W., Warren, H.B., and Kwiatkowski, D.J. (1999). Tsc2(+/-) mice develop tumors in multiple sites that express gelsolin and are influenced by genetic background. J. Clin. Invest. 104, 687–695. 10.1172/JCI7319.

38. Kobayashi, T., Minowa, O., Sugitani, Y., Takai, S., Mitani, H., Kobayashi, E., Noda, T., and Hino, O. (2001). A germ-line Tsc1 mutation causes tumor development and embryonic lethality that are similar, but not identical to, those caused by Tsc2 mutation in mice. Proc. Natl. Acad. Sci. U. S. A. 98, 8762–8767. 10.1073/pnas.151033798.

39. Guertin, D.A., Stevens, D.M., Thoreen, C.C., Burds, A.A., Kalaany, N.Y., Moffat, J., Brown, M., Fitzgerald, K.J., and Sabatini, D.M. (2006). Ablation in Mice of the mTORC Components raptor, rictor, or mLST8 Reveals that mTORC2 Is Required for Signaling to Akt-FOXO and PKCα, but Not S6K1. Dev. Cell 11, 859–871. 10.1016/j.devcel.2006.10.007.

40. Goorden, S.M.I., Hoogeveen-Westerveld, M., Cheng, C., Woerden, G.M. van, Mozaffari, M., Post, L., Duckers, H.J., Nellist, M., and Elgersma, Y. (2011). Rheb Is Essential for Murine Development. Mol. Cell. Biol. 31, 1672–1678. 10.1128/MCB.00985-10.

41. Kobayashi, T., Minowa, O., Kuno, J., Mitani, H., Hino, O., and Noda, T. (1999). Renal Carcinogenesis, Hepatic Hemangiomatosis, and Embryonic Lethality Caused by a Germ-Line Tsc2 Mutation in Mice 1.

42. Schrötter, S., Yuskaitis, C.J., MacArthur, M.R., Mitchell, S.J., Hosios, A.M., Osipovich, M., Torrence, M.E., Mitchell, J.R., Hoxhaj, G., Sahin, M., et al. (2022). The non-essential TSC complex component TBC1D7 restricts tissue mTORC1 signaling and brain and neuron growth. Cell Rep. 39. 10.1016/j.celrep.2022.110824.

43. Hernandez, O., Way, S., McKenna, J., and Gambello, M.J. (2007). Generation of a conditional disruption of the Tsc2 gene. genesis 45, 101–106. 10.1002/DVG.20271.

44. Shigeyama, Y., Kobayashi, T., Kido, Y., Hashimoto, N., Asahara, S.-I., Matsuda, T., Takeda, A., Inoue, T., Shibutani, Y., Koyanagi, M., et al. (2008). Biphasic response of pancreatic beta-cell mass to ablation of tuberous sclerosis complex 2 in mice. Mol. Cell. Biol. 28, 2971–2979. 10.1128/MCB.01695-07.

45. Yecies, J.L., Zhang, H.H., Menon, S., Liu, S., Yecies, D., Lipovsky, A.I., Gorgun, C., Kwiatkowski, D.J., Hotamisligil, G.S., Lee, C.H., et al. (2011). Akt stimulates hepatic SREBP1c and lipogenesis through parallel mTORC1-dependent and independent pathways. Cell Metab. 14, 21–32. 10.1016/j.cmet.2011.06.002.

46. Meikle, L., Talos, D.M., Onda, H., Pollizzi, K., Rotenberg, A., Sahin, M., Jensen, F.E., and Kwiatkowski, D.J. (2007). A Mouse Model of Tuberous Sclerosis: Neuronal Loss of Tsc1 Causes Dysplastic and Ectopic Neurons, Reduced Myelination, Seizure Activity, and Limited Survival. J. Neurosci. 27, 5546–5558. 10.1523/JNEUROSCI.5540-06.2007.

47. Uhlmann, E.J., Wong, M., Baldwin, R.L., Bajenaru, M.L., Onda, H., Kwiatkowski, D.J., Yamada, K., and Gutmann, D.H. (2002). Astrocyte-specific TSC1 conditional knockout mice exhibit abnormal neuronal organization and seizures. Ann. Neurol. 52, 285–296. 10.1002/ANA.10283.

48. Castets, P., and Rüegg, M.A. (2013). MTORC1 determines autophagy through ULK1 regulation in skeletal muscle. Autophagy 9, 1435–1437. 10.4161/AUTO.25722.

49. Yang, W., Pang, D., Chen, M., Du, C., Jia, L., Wang, L., He, Y., Jiang, W., Luo, L., Yu, Z., et al. (2021). Rheb mediates neuronal-activity-induced mitochondrial energetics through mTORC1-independent PDH activation. Dev. Cell 56, 811–825.e6. 10.1016/J.DEVCEL.2021.02.022/ATTACHMENT/447B028F-8163-4820-9762-CC77EC35230F/MMC2.PDF.

50. Ni, Q., Gu, Y., Xie, Y., Yin, Q., Zhang, H., Nie, A., Li, W., Wang, Y., Ning, G., Wang, W., et al. (2017). Raptor regulates functional maturation of murine beta cells. Nat. Commun. 2017 81 8, 1–13. 10.1038/ncomms15755.

51. Zou, J., Zhou, L., Du, X.X., Ji, Y., Xu, J., Tian, J., Jiang, W., Zou, Y., Yu, S., Gan, L., et al. (2011). Rheb1 Is Required for mTORC1 and Myelination in Postnatal Brain Development. Dev. Cell 20, 97–108. 10.1016/J.DEVCEL.2010.11.020/ATTACHMENT/774CC5FF-EF69-4B1F-8138-B45966FB4A35/MMC1.PDF.

52. Sengupta, S., Peterson, T.R., Laplante, M., Oh, S., and Sabatini, D.M. (2010). mTORC1 controls fasting-induced ketogenesis and its modulation by ageing. Nat. 2010 4687327 468, 1100–1104. 10.1038/nature09584.

53. Bentzinger, C.F., Romanino, K., Cloëtta, D., Lin, S., Mascarenhas, J.B., Oliveri, F., Xia, J., Casanova, E., Costa, C.F., Brink, M., et al. (2008). Skeletal Muscle-Specific Ablation of raptor, but Not of rictor, Causes Metabolic Changes and Results in Muscle Dystrophy. Cell Metab. 8, 411–424. 10.1016/j.cmet.2008.10.002.

54. Menon, S., Yecies, J.L., Zhang, H.H., Howell, J.J., Nicholatos, J., Harputlugil, E., Bronson, R.T., Kwiatkowski, D.J., and Manning, B.D. (2012). Chronic activation of mTOR complex 1 is sufficient to cause hepatocellular carcinoma in mice. Sci. Signal. 5, ra24. 10.1126/scisignal.2002739.

55. Guridi, M., Kupr, B., Romanino, K., Lin, S., Falcetta, D., Tintignac, L., and Rüegg, M.A. (2016). Alterations to mTORC1 signaling in the skeletal muscle differentially affect whole-body metabolism. Skelet. Muscle 6, 1–14. 10.1186/S13395-016-0084-8/FIGURES/6.

56. Sasaki, Y., Derudder, E., Hobeika, E., Pelanda, R., Reth, M., Rajewsky, K., and Schmidt-Supprian, M. (2006). Canonical NF-κB Activity, Dispensable for B Cell Development, Replaces BAFF-Receptor Signals and Promotes B Cell Proliferation upon Activation. Immunity 24, 729–739. 10.1016/j.immuni.2006.04.005.

57. Schwenk, F., Baron, U., and Rajewsky, K. (1995). A cre -transgenic mouse strain for the ubiquitous deletion of loxP -flanked gene segments including deletion in germ cells. Nucleic Acids Res. 23, 5080–5081. 10.1093/NAR/23.24.5080.

58. Dong, J., and Pan, D. (2004). Tsc2 is not a critical target of Akt during normal Drosophila development. Genes Dev. 18, 2479. 10.1101/GAD.1240504.

59. Schleich, S., and Teleman, A.A. (2009). Akt Phosphorylates Both Tsc1 and Tsc2 in Drosophila, but Neither Phosphorylation Is Required for Normal Animal Growth. PLoS One 4, e6305. 10.1371/JOURNAL.PONE.0006305.

60. Lai, Y.C., Kviklyte, S., Vertommen, D., Lantier, L., Foretz, M., Viollet, B., Hallén, S., and Rider, M.H. (2014). A small-molecule benzimidazole derivative that potently activates AMPK to increase glucose transport in skeletal muscle: comparison with effects of contraction and other AMPK activators. Biochem. J. 460, 363–375. 10.1042/BJ20131673.

61. Lemos, C., Schulze, V.K., Baumgart, S.J., Nevedomskaya, E., Heinrich, T., Lefranc, J., Bader, B., Christ, C.D., Briem, H., Kuhnke, L.P., et al. (2021). The potent AMPK inhibitor BAY-3827 shows strong efficacy in androgen-dependent prostate cancer models. Cell. Oncol. 44, 581–594. 10.1007/S13402-020-00584-8/FIGURES/5.

62. Cloëtta, D., Thomanetz, V., Baranek, C., Lustenberger, R.M., Lin, S., Oliveri, F., Atanasoski, S., and Rüegg, M.A. (2013). Inactivation of mTORC1 in the Developing Brain Causes Microcephaly and Affects Gliogenesis. J. Neurosci. 33, 7799–7810. 10.1523/JNEUROSCI.3294-12.2013.

63. Carson, R.P., Van Nielen, D.L., Winzenburger, P.A., and Ess, K.C. (2012). Neuronal and glia abnormalities in Tsc1-deficient forebrain and partial rescue by rapamycin. Neurobiol. Dis. 45, 369–380. 10.1016/J.NBD.2011.08.024.

64. Rolfe, D.F.S., and Brown, G.C. (1997). Cellular Energy Utilization and Molecular Origin of Standard Metabolic Rate in Mammals. REVIEWS 77.

65. Jaiswal, N., Gavin, M., Loro, E., Sostre-Colón, J., Roberson, P.A., Uehara, K., Rivera-Fuentes, N., Neinast, M., Arany, Z., Kimball, S.R., et al. (2022). AKT controls protein synthesis and oxidative metabolism via combined mTORC1 and FOXO1 signalling to govern muscle physiology. J. Cachexia. Sarcopenia Muscle 13, 495–514. 10.1002/JCSM.12846.

66. Xiong, X., Xie, R., Zhang, H., Gu, L., Xie, W., Cheng, M., Jian, Z., Kovacina, K., and Zhao, H. (2014). PRAS40 plays a pivotal role in protecting against stroke by linking the Akt and mTOR pathways. Neurobiol. Dis. 66, 43–52. 10.1016/J.NBD.2014.02.006.

67. Way, S.W., Mckenna Iii, J., Mietzsch, U., Reith, R.M., Cheng-Ju Wu, H., and Gambello, M.J. Loss of Tsc2 in radial glia models the brain pathology of tuberous sclerosis complex in the mouse. 10.1093/hmg/ddp025.

68. Magri, L., Cambiaghi, M., Cominelli, M., Alfaro-Cervello, C., Cursi, M., Pala, M., Bulfone, A., Garca-Verdugo, J.M., Leocani, L., Minicucci, F., et al. (2011). Sustained activation of mTOR pathway in embryonic neural stem cells leads to development of tuberous sclerosis complex-associated lesions. Cell Stem Cell 9, 447–462. 10.1016/j.stem.2011.09.008.

69. Risson, V., Mazelin, L., Roceri, M., Sanchez, H., Moncollin, V., Corneloup, C., Richard-Bulteau, H., Vignaud, A., Baas, D., Defour, A., et al. (2009). Muscle inactivation of mTOR causes metabolic and dystrophin defects leading to severe myopathy. J. Cell Biol. 187, 859–874. 10.1083/JCB.200903131.

70. Baraldo, M., Zorzato, S., Dondjang, A.H.T., Geremia, A., Nogara, L., Dumitras, A.G., Canato, M., Marcucci, L., Nolte, H., and Blaauw, B. (2022). Inducible deletion of raptor and mTOR from adult skeletal muscle impairs muscle contractility and relaxation. J. Physiol. 600, 5055–5075. 10.1113/JP283686.

71. Padi, S.K.R., Singh, N., Bearss, J.J., Olive, V., Song, J.H., Cardó-Vila, M., Kraft, A.S., and Okumura, K. (2019). Phosphorylation of DEPDC5, a component of the GATOR1 complex, releases inhibition of mTORC1 and promotes tumor growth. Proc. Natl. Acad. Sci. U. S. A. 116, 20505–20510. 10.1073/PNAS.1904774116/SUPPL_FILE/PNAS.1904774116.SAPP.PDF.

72. Haas, J.T., Miao, J., Chanda, D., Wang, Y., Zhao, E., Haas, M.E., Hirschey, M., Vaitheesvaran, B., Farese, R. V., Kurland, I.J., et al. (2012). Hepatic insulin signaling is required for obesity-dependent expression of SREBP-1c mRNA but not for feeding-dependent expression. Cell Metab. 15, 873–884. 10.1016/j.cmet.2012.05.002.

73. de la Calle Arregui, C., Plata-Gómez, A.B., Deleyto-Seldas, N., García, F., Ortega-Molina, A., Abril-Garrido, J., Rodriguez, E., Nemazanyy, I., Tribouillard, L., de Martino, A., et al. (2021). Limited survival and impaired hepatic fasting metabolism in mice with constitutive Rag GTPase signaling. Nat. Commun. 2021 121 12, 1–20. 10.1038/s41467-021-23857-8.

74. Trefts, E., Gannon, M., and Wasserman, D.H. (2017). The liver. Curr. Biol. 27, R1147– R1151. 10.1016/j.cub.2017.09.019.

75. Cangelosi, A.L., Puszynska, A.M., Roberts, J.M., Armani, A., Nguyen, T.P., Spinelli, J.B., Kunchok, T., Wang, B., Chan, S.H., Lewis, C.A., et al. (2022). Zonated leucine sensing by Sestrin-mTORC1 in the liver controls the response to dietary leucine. Science 377, 47. 10.1126/SCIENCE.ABI9547.

76. Hee Um, S., Frigerio, F., Watanabe, M., dé ric Picard, F., Joaquin, M., Sticker, M., Fumagalli, S., Allegrini, P.R., Kozma, S.C., Auwerx, J., et al. (2004). Absence of S6K1 protects against age-and diet-induced obesity while enhancing insulin sensitivity.

77. Khamzina, L., Veilleux, A., Bergeron, S., and Marette, A. (2005). Increased Activation of the Mammalian Target of Rapamycin Pathway in Liver and Skeletal Muscle of Obese Rats: Possible Involvement in Obesity-Linked Insulin Resistance. 10.1210/en.2004-0921.

78. Mannick, J.B., and Lamming, D.W. (2023). Targeting the biology of aging with mTOR inhibitors. Nat. Aging 2023 36 3, 642–660. 10.1038/s43587-023-00416-y.

79. Zhang, H.H., Huang, J., Düvel, K., Boback, B., Wu, S., Squillance, R.M., Wu, C.L., and Manning, B.D. (2009). Insulin stimulates adipogenesis through the Akt-TSC2-mTORC1 pathway. PLoS One. 10.1371/journal.pone.0006189.

80. Durkin, M., Qian, X., Popescu, N., and Lowy, D. (2013). Isolation of Mouse Embryo Fibroblasts. BIO-PROTOCOL 3. 10.21769/BIOPROTOC.908.

81. Byles, V., Cormerais, Y., Kalafut, K., Barrera, V., Hughes Hallett, J.E., Sui, S.H., Asara, J.M., Adams, C.M., Hoxhaj, G., Ben-Sahra, I., et al. (2021). Hepatic mTORC1 signaling activates ATF4 as part of its metabolic response to feeding and insulin. Mol. Metab. 53, 101309. 10.1016/J.MOLMET.2021.101309.

82. Wright, A., Hall, A., Daly, T., Fontelonga, T., Potter, S., Schafer, C., Lindsley, A., Hung, C., Bodamer, O., and Gussoni, E. (2021). Lysine methyltransferase 2D regulates muscle fiber size and muscle cell differentiation. FASEB J. 35, e21955. 10.1096/FJ.202100823R.

83. Wu, Y.F., Lapp, S., Dvoretskiy, S., Garcia, G., Kim, M., Tannehill, A., Daniels, L., and Boppart, M.D. (2022). Optimization of a pericyte therapy to improve muscle recovery after limb immobilization. J. Appl. Physiol. 132, 1020–1030. 10.1152/JAPPLPHYSIOL.00700.2021/ASSET/IMAGES/LARGE/JAPPLPHYSIOL.00700.2021_F004.JPEG.

84. Esper, M.E., Kodippili, K., and Rudnicki, M.A. (2023). Immunofluorescence Labeling of Skeletal Muscle in Development, Regeneration, and Disease. Methods Mol. Biol. 2566, 113–132. 10.1007/978-1-0716-2675-7_9.

85. Schmidt, E.K., Clavarino, G., Ceppi, M., and Pierre, P. (2009). SUnSET, a nonradioactive method to monitor protein synthesis. Nat. Methods 2009 64 6, 275–277. 10.1038/nmeth.1314.

