## Supplementary figures 1-4 for "AKT-mediated phosphorylation of TSC2 controls stimulus- and tissue-specific mTORC1 signaling and organ growth"

### Supplementary figure legends

#### Figure S1: Data supporting Figure 1.

(A) Representative immunoblots of lysates from primary C57/BL6J hepatocytes, serum-starved for 16 h followed by a 15-min stimulation with insulin (100 nM) after a 30-min pretreatment with vehicle (-), MK2206 (2 µM, AKTi), trametinib (5 µM, MEKi), both, or rapamycin (20nM, Rap), as indicated.

(B) Representative immunoblots of lysates from primary C57/BL6J myotubes, serum-starved for 6 h followed by a 15-min stimulation with insulin (100 nM) after a 30-min pretreatment with vehicle (-), MK2206 (2 µM, AKTi), trametinib (5 µM, MEKi), both, or Rapamycin (20nM, Rap), as indicated.

(C) Representative immunoblots of lysates from primary C57/BL6J cortical neurons following 4 h of growth factor and amino acid withdrawal, followed by a 15-min stimulation with IGF1 (100 ng/mL) after a 30-min pretreatment with vehicle (-), MK2206 (2 µM, AKTi), trametinib (5 µM, MEKi), both, or rapamycin (20nM, Rap), as indicated.

(D) Schematic of the *Rosa26* locus (R26) with the Flag-TSC2 targeting vector inserted. Locations of the different primers used in (E-G) and listed in the methods are indicated.

(E) PCR results with the indicated primers from (D) showing homologous recombination of the TSC2-WT and -5A targeting vectors with the *Rosa26* locus in 15.5 dpc whole embryos. DNA isolated from a C57BL/6J (B6) embryo was used as a negative control.

(F) PCR results with the indicated primers from (D) showing presence (left) or CRE-mediated deletion (right) of the Lox-Stop-Lox cassette in *Rosa26-LSL-TSC2-WT* or *-5A;* $Tsc2^{fl/fl}$ (Cre -) and *Rosa26 -TSC2-WT* or *-5A;* $Tsc2^{-/-}$ (Cre +) 15.5 dpc whole embryos.

(G) Representative PCR genotyping results of the *Rosa26* locus (left) with TSC2-WT or -5A transgene insertions and the mouse $Tsc2^{fl/fl}$ locus (right) before (mTsc2 Flox) or after Cre-mediated deletion (Δ mTSC2) in 15.5 dpc whole embryos. Note, the TSC2-5A transgene lacks an SpeI site (specified in D) present in the TSC2-WT transgene.

(H) Representative immunoblots demonstrating CRE-mediated expression of the Flag-tagged human TSC2-WT and -5A and GFP proteins in extracts from *Rosa26-LSL-TSC2-WT* or *-5A;* $Tsc2^{fl/fl}$ (Cre -) and *Rosa26-TSC2-WT* or *-5A;* $Tsc2^{-/-}$ (Cre +) 15.5 dpc whole embryos.

#### Figure S2: Data supporting Figure 2.

(A) Representative immunoblots of lysates from primary TSC2-WT and TSC2-5A MEFs serum-starved for 16 h and then treated for 30 min with vehicle (-), MK2206 (2 µM) or trametinib (5 µM), as indicated.

(B) Quantification of pS6K1/S6K1 ratio from (A) graphed as mean percentage of vehicle-treated TSC2-WT MEFs ± SEM; N = 3 biological replicates per genotype.

Statistical analysis by ordinary two-way ANOVA with Tukey's multiple comparisons test, $*p < 0.05$, $**p < 0.01$, $***p < 0.001$.

**Figure S3: Data supporting Figure 3.**

(A) Male organ weights, normalized (bottom) or not (top) to body weight (BW) in 9-week-old TSC2-5A mice, graphed as mean percentage of TSC2-WT littermates ± SEM; N = 7 TSC2-WT and 7 TSC2-5A.

(B) Non-normalized male organ weights in 16-week-old TSC2-5A mice, graphed as mean percentage of TSC2-WT littermates ± SEM; N = 6 TSC2-WT and 8 TSC2-5A.

(C) Female organ weights, normalized (bottom) or not (top) to body weight (BW) in 9-week-old TSC2-5A mice, graphed as mean percentage of TSC2-WT littermates ± SEM; N = 7 TSC2-WT and 7 TSC2-5A

(D) Immunoblots of brain lysates from ad libitum fed 9-week-old male TSC2-WT and TSC2-5A littermates. Data quantified in Figure 3G. N = 7 TSC2-WT and 7 TSC2-5A.

Statistical analysis (A-C) by Mann-Whitney test, $*p < 0.05$.

**Figure S4: Data supporting Figure 4.**

(A) Immunoblots of skeletal muscle and liver lysates from ad libitum fed 9-week-old male TSC2-WT and TSC2-5A littermates. Data quantified in Figure 4A. N = 7 TSC2-WT and 7 TSC2-5A.

(B) Echo-MRI fat mass measurements of male littermates at 3, 6, and 12 months, graphed as mean ± SEM; N = 12 TSC2-WT and 9 TSC2-5A.

(C) Echo-MRI lean and fat mass measurements of female littermates at 3 and 12 months, graphed as mean ± SEM; N = 10 TSC2-WT and 7 TSC2-5A.

Statistical analysis (B, C) by ordinary two-way ANOVA with Tukey's multiple comparisons test, $*p < 0.05$, $**p < 0.01$.

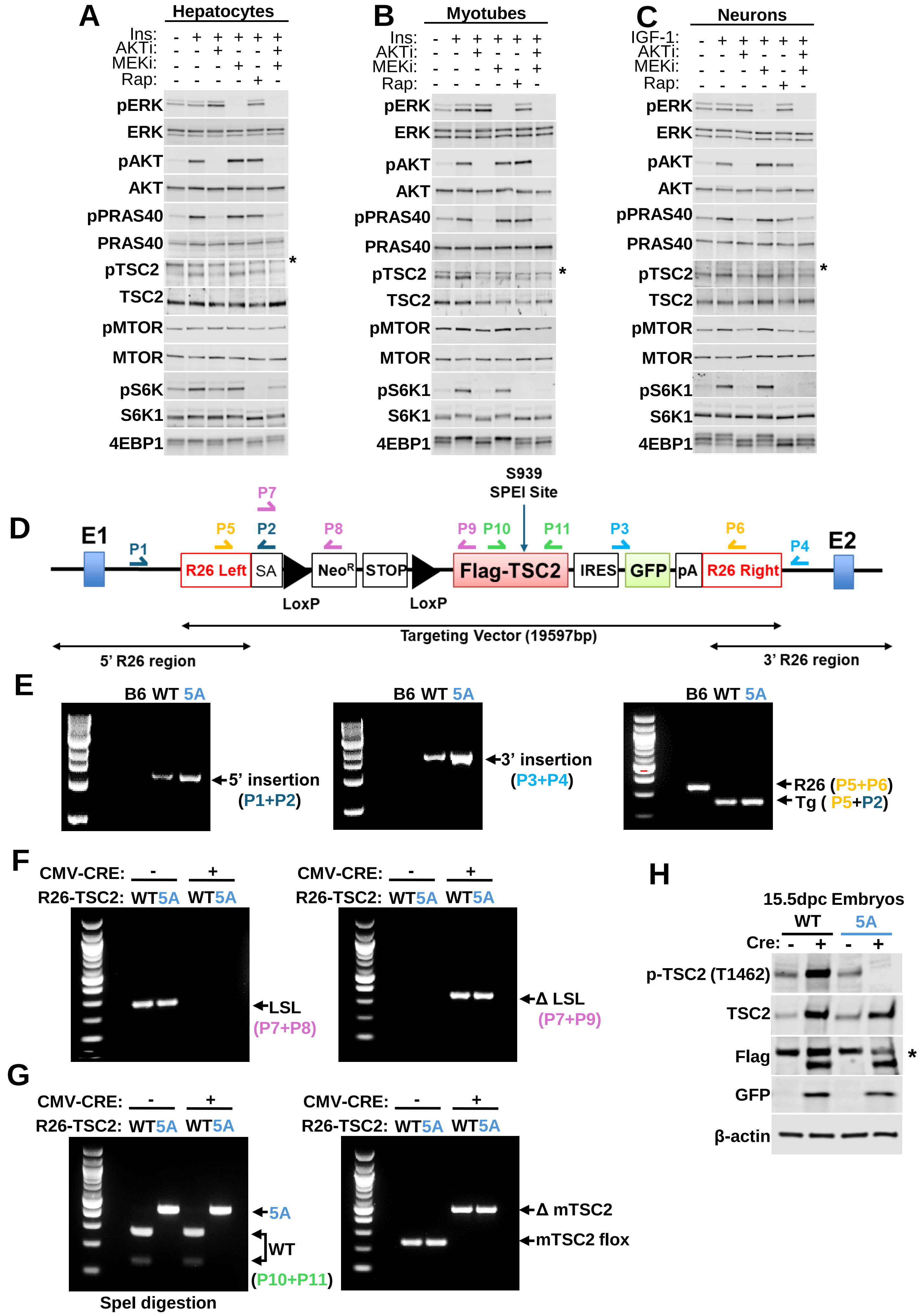

A
Hepatocytes
Ins:
AKTi:
MEKi:
Rap:
pERK
ERK
pAKT
AKT
pPRAS40
PRAS40
pTSC2
TSC2
pMTOR
MTOR
pS6K
S6K1
4EBP1
B
Myotubes
pS6K1
C
Neurons
IGF-1:
D
E1
P1
P5
P2
P7
P8
S939
SPEI Site
P9
P10
P11
P3
P6
P4
E2
R26 Left
SA
NeoR
STOP
Flag-TSC2
IRES
GFP
pA
R26 Right
LoxP
LoxP
Targeting Vector (19597bp)
5' R26 region
3' R26 region
E
B6 WT 5A
5' insertion
(P1+P2)
3' insertion
(P3+P4)
R26 (P5+P6)
Tg ( P5+P2)
F
CMV-CRE: - +
R26-TSC2: WT5A WT5A
LSL
(P7+P8)
Δ LSL
(P7+P9)
G
5A
WT
(P10+P11)
Δ mTSC2
mTSC2 flox
SpeI digestion
H
15.5dpc Embryos
WT
5A
Cre: - + - +
p-TSC2 (T1462)
TSC2
Flag
GFP
β-actin


Fig-S1

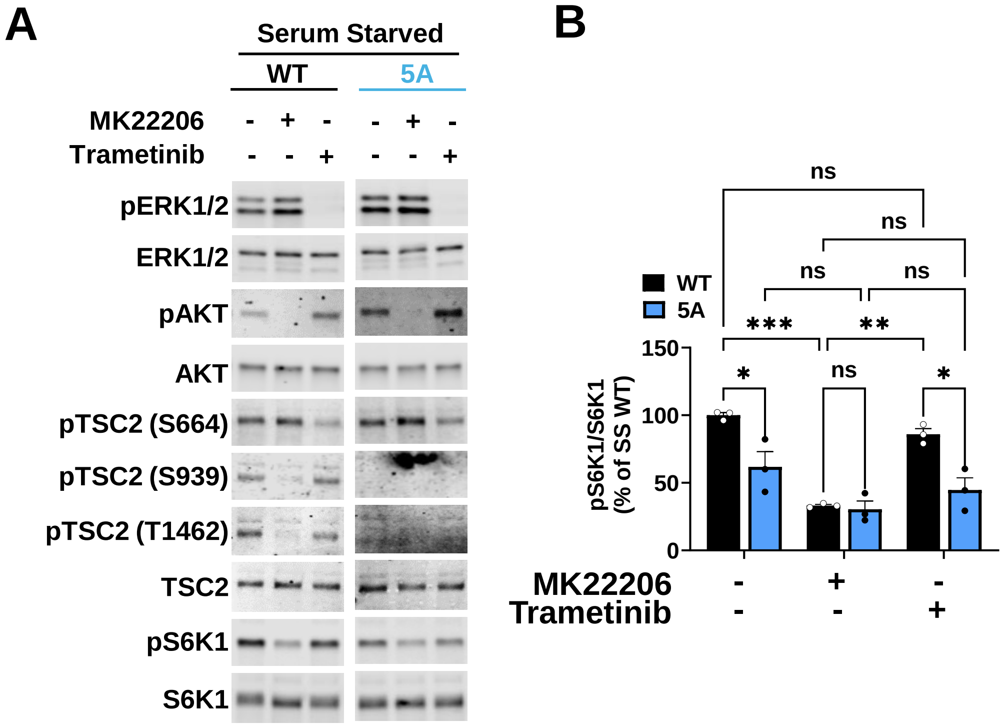
A
Serum Starved
WT
5A
MK22206 - + - - + -
Trametinib - - + - - +
pERK1/2
ERK1/2
pAKT
AKT
pTSC2 (S664)
pTSC2 (S939)
pTSC2 (T1462)
TSC2
pS6K1
S6K1
B
WT
5A
ns
***
**
*
pS6K1/S6K1
(% of SS WT)
150
100
50
0
MK22206 - + -
Trametinib - - +

Fig-S2

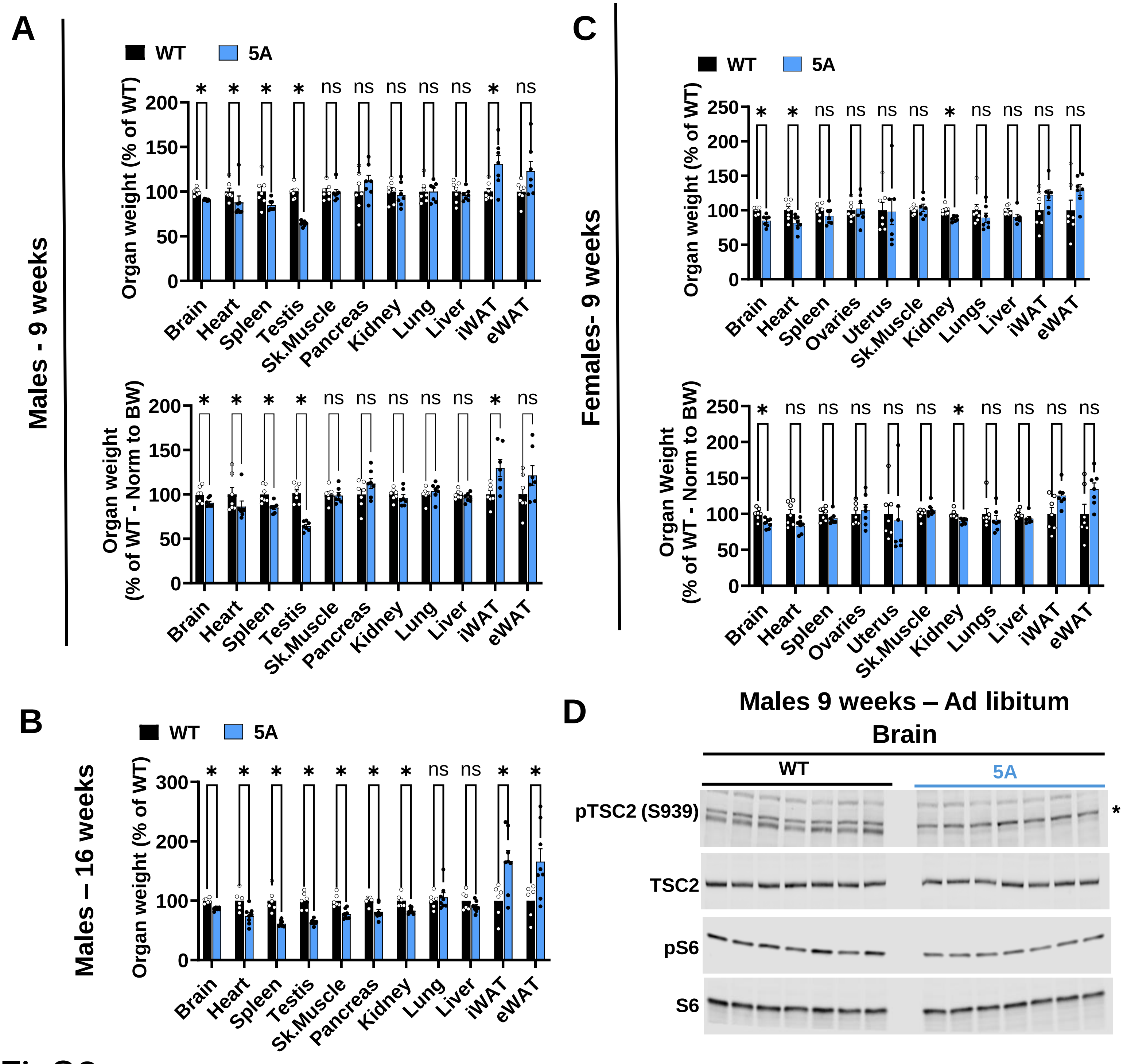
A
Males - 9 weeks
WT
5A
Organ weight (% of WT)
Brain
Heart
Spleen
Testis
Sk.Muscle
Pancreas
Kidney
Lung
Liver
iWAT
eWAT
Organ weight (% of WT - Norm to BW)
B
Males – 16 weeks
C
Females- 9 weeks
Ovaries
Uterus
Lungs
Organ Weight (% of WT - Norm to BW)
D
Males 9 weeks – Ad libitum
Brain
pTSC2 (S939)
TSC2
pS6
S6

FigS3

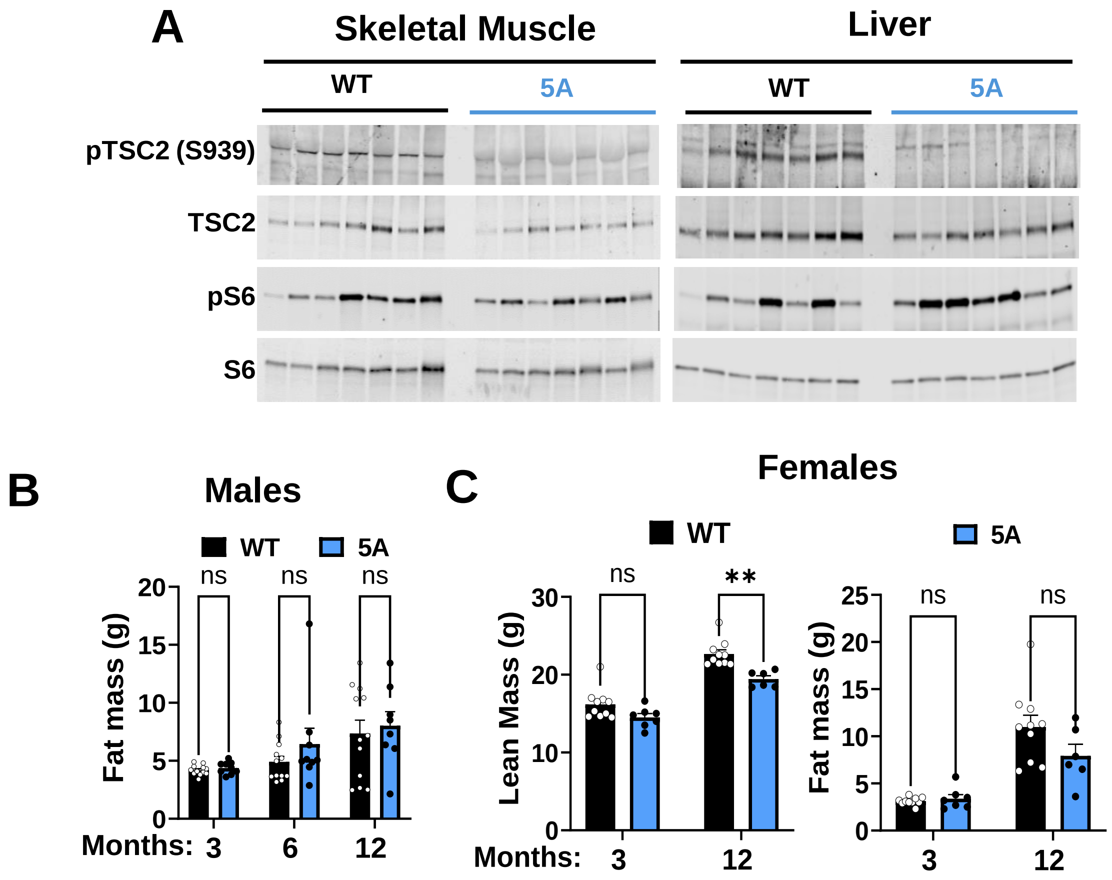
A
Skeletal Muscle
Liver
WT
5A
WT
5A
pTSC2 (S939)
TSC2
pS6
S6
B
Males
WT
5A
ns
ns
ns
Fat mass (g)
Months: 3
6
12
C
Females
WT
5A
ns
**
Lean Mass (g)
Months: 3
12
ns
ns
Fat mass (g)
3
12

FigS4
